# The phosphoproteome is a first responder in tiered cellular adaptation to chemical stress followed by proteomics and transcriptomics alteration

**DOI:** 10.1101/2022.04.07.487458

**Authors:** Peiru Chen, Yuan Li, Feng Xu, Zhenpeng Zhang, Tao Zuo, Jiabin Guo, Kaixuan Li, Shu Liu, Suzhen Li, Jian Yin, Lei Chang, Predrag Kukic, Mark Liddell, Liz Tulum, Paul Carmichael, Shuangqing Peng, Jin Li, Qiang Zhang, Ping Xu

## Abstract

Next-generation risk assessment for environmental chemicals and ingredients in consumer products involves a weight of evidence (WoE) framework integrating a suite of new approach methodologies (NAMs) based on points of departure (PoD) obtained from *in vitro* assays. Omics techniques provide broad coverages of the molecular toxicity pathway space. Transcriptomics assays especially play a leading role by providing relatively conservative PoDs in comparison with apical endpoints. However, it is unclear whether and how parameters measured using other omics technicquesparticipate in the cellular response to chemical perturbations, especially at exposure levels below the transcriptomically defined PoD. Multi-omics coverage may provide additional sensitive or confirmative biomarkers to complement and reduce the uncertainty in safety decisions made using targeted and transcriptomics assays. In the present study, we compared changes in transcriptomics, proteomics and phosphoproteomics with two prototype compounds, coumarin, as a main study and doxorubicin, as a complementary study to understand the sensitivity of the three omics techniques in response to chemically-induced changes in HepG2 and AC16 cells. With measurements obtained for multiple chemical concentrations and time points, we demonstrated that, compared with proteomics and transcriptomics, phosphoproteomics alterations occur not only earlier in time as expected, but also at much lower chemical concentrations and hence are proximal to the very early effects induced by chemical exposure. The phosphoproteomics changes appear to approach maximum when the transcriptomics alterations begin to be initiated. The results are consistent with a tiered framework of cellular response to chemical insults, where posttranslational modification of preexisting proteins is first seen before transcriptomics induction is engaged to launch a more energy-expensive defense that defines a useful PoD. We conclude that as the cost becomes more affordable, proteomics covering posttranslational modifications can be utilized to provide a more complete coverage of chemical-induced cellular alteration and supplement transcriptomics-based health safety decision making.

## Introduction

Next-generation chemical risk assessment (NGRA) is moving steadily toward relying on cell- or organoid-based assays to identify useful *in vitro* points of departure (PoD) that can be used in safety decision-making^1–3^. In assayed cells, a myriad of molecular and morphological changes can be induced by chemicals, depending on the physiochemical property of the chemicals, their concentrations and exposure durations. Some chemicals can induce these changes through recognizing specific molecular targets in the cells, such as receptors or enzymes, to activate or inhibit one or multiple toxicity pathways. Alternatively, reactive chemicals, which do not necessarily have a specific molecular target, can induce cellular responses via activating stress pathways^4, 5^. Regardless of the nature of chemical actions, many of these induced changes can be measured economically and used as biomarkers to cover a variety of specific cellular and molecular endpoints, such as receptor binding, enzyme inhibition, immunomodulation, DNA damage, stress responses, and mitochondrial activity^4, 6–10^.

Targeted analyses focusing on a suite of specific cellular and molecular endpoints cannot provide sufficient coverage of the biological space perturbed by a chemical, especially for novel chemicals whose mode of actions are unknown. Yet, any apical endpoint change indicative of impaired health must be underpinned by some alterations at the omics levels, including the metabolome, proteome, transcriptome, epigenome and genome^11–13^. Technical advancements of omics tools in the past two decades have enabled global coverage of molecular changes occurring in cells that is unmatched by traditional targeted analyses. Due in a large part to the relatively lower cost, broader biological coverage, higher S/N ratio, and smaller sample size, transcriptomics has been at the top of the omics assay list in the past decades for examining cellular response to biological and chemical perturbations. Starting with early functional toxicogenomics studies using gene microarrays and followed by next-generation RNA sequencing in recent years, transcriptomics has become a routine assay in toxicological research *in vivo* and *in vitro*^14–16^. An increasing number of studies have revealed a consensus that seems to hold across the chemical and biological space – the PoD as defined by the most sensitive transcriptomics pathway alterations in short-term animal or *in vitro* cell assays appears to be within a similar order of magnitude to the doses or concentrations causing pathological outcomes in long-term animal studies for many legacy chemicals whose toxicities are known^17–23^. This concordance in PoD between transcriptional and apical endpoint changes is largely conserved for both cancer and noncancer endpoints^18^. Thus, transcriptomics is emerging as one of the most promising new approach methodologies (NAMs) with broad coverage for chemical toxicity screening. As a result of these encouraging developments in toxicogenomics, developing high-throughput transcriptomics *in vitro* assays and associated bioinformatic tools is one of the top priorities in the next-decade blueprint recently launched by the US EPA^10^.

It is becoming general practice now to use the beginning of transcriptomics alterations, often defined by the benchmark dose (BMD) either at gene or pathway levels^24, 25^, to provide a PoD for safety assessment. The cutoff values for the number of genes or fold-change of differential expression selected to determine the BMD may introduce uncertainty, along with uncertainties associated with determining the BMD confidence lower limit. Beside the statistical considerations, mechanistically speaking, it is not completely clear how transcriptomics alterations are linked to adverse apical endpoint changes for broad classes of chemicals. Receptor-mediated or target-specific transcriptomics alterations, when of sufficient magnitude and duration, can be led to an alternative or compromised functional state in a cell, thus leading to potential apical adversity. In comparison, reactive chemicals can elicit nonspecific adaptive cellular stress responses requiring transcriptional induction of key stress genes^4^. Cellular stress responses are bioenergetically demanding as a suite of stress proteins need to be synthesized *de nova* in large quantities^26–31^. As a result, cells may enter a survival mode where they undergo global translational and metabolic reprogramming to preserve and divert energy for the stress response while specialized cell functions may be compromised^32, 33^. Therefore, the onset of adaptive transcriptional stress control, for the purpose of cell survival, may be regarded as the onset of functional adversity.

If apical adverse outcomes involve obligatory transcriptome-mediated responses, it begs the question ‘what are the mechanisms that protect cells from the chemical insult and preserve functions under exposures below the transcriptomically-defined PoD doses?’. If known, these non-transcriptomics molecular changes, occurring at lower doses, may be exploited to develop sensitive biomarkers that can help reduce the uncertainty associated with PoD in a WoE safety assessment framework. Many posttranscriptional control mechanisms appear to be engaged by activating pre-existing stress proteins that are not activated at nonstressed conditions^5, 34^. This activation can be mediated through posttranslational modifications such as phosphorylation, oxidation, and acetylation^35–37^. Compared with the resource-heavy transcriptional adaptation, these posttranslational processes are fast-responding and demand much less energy. Thus cells are likely to cope with chemical stresses in a two-tiered manner^5^. At low concentrations, cells use posttranslational regulation of pre-existing stress proteins to maintain homeostasis without affecting their specialized functions as no global translational and metabolic reprogramming is engaged. At higher concentrations, pre-existing stress proteins are activated and transcriptional induction is initiated to increase their abundance to continue to maintain homeostasis and survival.

Therefore, the transition from posttranslational to transcriptional control, where proteomics changes are fully engaged and transcriptomics induction is initiated, may mark a PoD (i.e., the transcriptomics PoD), where specialized cell functions begin to be compromised resulting in reduced fitness. In cases where there are typically low exposures, such as environmental exposure or in the situation of cosmetic usage of ingredients at low concentrations, posttranslational-mediated protection may be at play. At higher exposures, transcriptomically-mediated protection may have collateral adverse consequence, leading to safety concerns. As a result of the two-tiered stress response, it is worth monitoring posttranslational changes as additional biomarkers that can more sensitively detect and corroborate the PoD that is defined by the onset of transcriptomics and functional alterations.

In the present study, we aim to compare cellular responses at multi-omics levels, including transcriptomics, proteomics and phosphoproteomics to systemically test the two-tiered response hypothesis on a global scale and ascertain whether phosphoproteomics is more sensitive in response to chemical challenges. coumarin and doxorubicin were selected as our case study chemicals. They represent a safe compound widely used in cosmetics and a pharmaceutical with known cardiotoxicity, respectively, for which historical *in vitro* data generated in HepG2 cells, a hepatocyte cell line, and AC16 cells, a human cardiomyocyte cell line exist^8, 9^. In the present study, we conducted multi-concentration and multi-time point transcriptomics, proteomics and phosphoproteomics studies in HepG2 and AC16 cells with these two chemicals. Our results demonstrated that both concentration-wise and time-wise, phosphoproteomics is much more sensitive in detecting cellular responses to chemical exposures than proteomics and transcriptomics; phosphoproteomics provides unique values to an integrated transcriptomics-centered multi-omics approach for toxicity testing.

## Materials and Methods

### Reagents and cell strains

Phospho-(Ser/Thr) Phe Antibody (9631S) was purchased from Cell Signaling Technology (Danvers, MA, USA). Pan Phospho-Tyrosine Rabbit pAb (AP0905) was obtained from ABclonal (Wuhan, Hubei, China). Antibodies against GAPDH (CW0100) goat anti rabbit IgG-HRP (CW0103S) and goat anti-mouse IgG-HRP (CW0102) were from CoWin Biosciences (Taizhou, Jiangsu, China). Acetylated trypsin ^38^ were from Enzyme & Spectrum (Beijing, China). Coumarin (C4261-50G), phosphatase inhibitor cocktail 2 (P5726) and 3 (P0044) were purchased from Sigma-Aldrich (St. Louis, MO, USA), and Doxorubicin (Adriamycin) HCl (NSC 123127) was from Selleck (Houston, Texas, USA). EDTA-free protease inhibitor cocktail was from Roche (Basel, Basel-Stadt, Switzerland). HepG2 was obtained from ATCC (American type culture collection, Washington D.C., USA).

### Cell culture and treatment

For the treatment with coumarin, HepG2 cells were seeded into a 10-cm petri dish containing Dulbecco’s Modified Eagle Medium (DMEM) supplemented with 10% fetal bovine serum (FBS), 100 U/mL penicillin and 100 mg/mL streptomycin at 37°C in a humidified 5% CO_2_ atmosphere incubator. Serially diluted chemicals were added on the following day without or with varied concentrations (0.001, 0.01, 0.1, 1, 10, and 100 μM) of coumarin at a 10-fold increment, which didn’t show any detectable changes in cell count and viability in our previous study^39^. All coumarin experiments were performed in triplicate. Cells were recovered after treatment for 6 hours and 24 hours for transcriptomics analysis, and after treatment for 10 min and 24 h for proteomics and phosphoproteomics analysis. For the treatment with doxorubicin, AC16 cells, were cultured using the same conditions as for HepG2 cells. AC16 cells were treated with 10 nM doxorubicin for 10 min, 30 min and 6 hours before recovery for transcriptomics, proteomics and phosphoproteomics analysis.

### Transcriptomics and data processing

Approximately 1×10^7^ fresh cells as described above were washed twice with ice-cold PBS then scraped with PBS and centrifuged at 12,000 g for 10 min at 4 °C. Total RNA was extracted using TRIzol reagent (Invitrogen, Carlsbad, CA, USA) and treated with RNase-free DNase I (Promega, Madison, WI, USA). The concentration of RNA was measured by a NanoPhotometer spectrophotometer and the quality of extracted RNA was checked with standard electrophoresis for RNA as previously described^40^. RNA was randomly fragmented, and then reversely transcribed to synthesize the first strand of cDNA using SmartScribe Rtase (Clontech, Mountain View, CA, USA). Owing to the terminal transferase activity, (dC) 3 was added to the 5’-terminus during synthesis of the first strand of cDNA. The second strand was synthesized using the adapter 1 primer containing (dG) 3 at the 5’-terminus. cDNAs sized 400–500 bp were isolated from 2% agarose gels after electrophoresis to generate the libraries. The libraries were further amplified by PCR with Pfu polymerase (NEB, Beverly, MA, USA). All libraries were sequenced with the adapter 1 primer using the Illumina HiSeq-2500. Cleaned reads were aligned to the Ensemble GRCh38 reference genome with Hisat2 using strand specific parameters.

The value of fragments per kilobase of transcript per million mapped reads (FPKM) was used to clean the data and for data quality control. We set a stringent filtering criterion of FPKM value > 1.0 in at least one out of all 42 samples to reduce false positives. After the genes were filtered, the expressed genes and transcripts were assembled and quantified by running StringTie^41^. For HepG2 cells with coumarin treatment, the differentially expressed genes for the RNA-seq data were calculated with gene count value by using edgeR package. Each concentration condition was compared with vehicle control. A criterion of ratio > 1.5 or < 0.67 with p value < 0.05 was used as the threshold for differentially expressed genes.

### Protein extraction

Approximately 5×10^7^ fresh cells were washed with ice-cold PBS. Cell lysis was performed on ice and sonicated for 4 min (2-s on and 4-s off, amplitude 25%) in buffer containing 9 M Urea, 10 mM Tris–HCl (pH 8.0), 30 mM NaCl, 5 mM IAA, 5 mM Na4P2O7, 100 mM Na2HPO4 (pH 8.0), 1 mM NaF, 1 mM Na3VO4, 1 mM Sodium glycerophophate, 1% phosphatase inhibitor cocktail 2 & 3 (Sigma) and protease inhibitor cocktail. Unbroken debris was eliminated by centrifugation (14,800 g) at 4 °C for 15 min, and the supernatant was collected and quantified by a gel-assisted method as previously described ^42^.

### Protein digestion by gel-aided strategy

Extracted protein samples (1 mg) of equal amount for all cells were reduced with 10 mM of dithiothreitol (DTT) at 45 °C for 30 min, alkylated with 20 mM of IAA in the dark at room temperature (RT) for 30 min, and cleaned by a short 10% SDS-PAGE. Each gel lane was cut into approximately 1 mm^3^ small cubes and digested with Ac-trypsin at 37 °C for 14 h to generate short peptides ^38, 43^. The tryptic peptides were collected with extracting buffer (5% formic acid (FA), 50% acetonitrile (ACN)) and dried with a vacuum dryer. About 5% of tryptic peptides were used for proteomics profiling experiments and the remaining 95% for phosphoproteomics study.

### Phosphopeptide enrichment and fractionation

Tryptic peptides were desalted by reverse-phase C18 Sep-Pak extraction cartridges (Waters Corporation, Milford, MA, USA). Cleaned peptides were resuspended in buffer A (1% ACN, 0.1% trifluoroacetic acid) and separated into 12 fractions through a Bonna-Agela C18 column (5 µm reverse-phase fused-silica, 4.6 × 250 mm column) on a RIGOL-L3000 HPLC (RIGOL, Beijing, China).The phosphor-peptides in each fraction were enriched with a Ti^4+^-immobilized metal ion affinity chromatography (IMAC) method as previously described with slight modification^44, 45^. Briefly, dried peptides were dissolved in 500 μL binding buffer (80% ACN, 6% trifluoroacetic acid (TFA) in ddH_2_O) and the same volume of loading buffer (10% 500 mM NH_4_HCO_3_, 5% ACN in ddH_2_O) followed by addition of 25 mg Ti^4+^-IMAC beads. Then, the mixed samples were vortexed at RT for 30 minutes, and centrifuged at 13,000 g for 6 min. The supernatant was removed, washed with washing buffer 1 (50% ACN, 6% TFA in 200 mM NaCl) once and washing buffer 2 (30% ACN, 0.1% TFA in ddH_2_O) twice for 30 min. After removing the supernatants, the phosphopeptides were eluted with elution buffer (10% NH3•H2O) for 15 min, and then sonicated in ice water for another 15 min. The mixtures were centrifuged at 17,000 g for 6 min, and the supernatant was transferred into a new tube and immediately acidified with 5% FA and lyophilized before LC-MS analysis.

### LC-MS/MS

#### Total cell lysate profiling

The enriched phoshopeptides were separated with self-packed StageTip column^46^. Tryptic peptides were resuspended in loading buffer (0.1% formic acid (FA) in ddH2O) and analyzed using an EASY-nLC 1000 ultra-performance liquid chromatography (UPLC) system (Thermo Fisher Scientific, San Jose, CA, USA) equipped with a self-packed capillary column (75 μm i.d. × 15 cm, 3 μm C18 reverse-phase fused-silica), with an 80 min nonlinear gradient at a flow rate of 600 nL/min. The gradient was composed of an increase from 5% to 12% solvent B (0.1% FA in 98% ACN) for 8 min, 12% to 24% in 46 min, 24% to 36% in 20 min, 36% to 95% in 1 min, and a final hold at 95% for the last 5 min.

The eluted peptides were ionized under high voltage (2 kV) and analyzed online using a Lumos^TM^ mass spectrometer (Thermo Fisher Scientific, San Jose, CA, USA). The initial MS spectrum (MS_1_) was analyzed over a mass range of 300−1400 Da with a resolution of 60,000 at m/z 200. The automatic gain control (AGC) was set to 5 × 10^5^, and the maximum injection time (MIT) was 50 ms. Subsequent MS/MS spectrum (MS_2_) was analyzed by using a data-dependent acquisition mode to search for the most intense ions within 3 s, which were detected in the Orbitrap at a resolution of 15,000 after fragmentation. For each scan, the threshold for triggering MS_2_ was set at 5,000 counts, isolation width was 1.6 m/z, normalized collision energy (NCE) was 35%, AGC was 5 × 10^4^, and MIT was 60 ms. Precursor ion charge-state screening was enabled and all unassigned charge states, as well as singly charged species, were rejected. The dynamic exclusion was set at 25 s to avoid repeated detection of the same precursor ions.

#### Phospho-peptide profiling

Phospho-peptide samples were dissolved in loading buffer (0.1% formic acid (FA) in ddH_2_O) and injected onto a UPLC (Ultimate 3000, Thermo Fisher Scientific, San Jose, CA, USA) equipped with self-packed capillary column (150 μm i.d.×12 cm, 1.9 μm C18 reverse-phase fused-silica). The peptides were eluted with a 78-min nonlinear gradient at a flow rate of 600 nL/min with an elution gradient and eluted samples were analyzed by Q-Exactive HF or LUMOS (Thermo Fisher Scientific, San Jose, CA, USA) in a data-dependent acquisition (DDA) mode. Briefly, the spray voltage was set as 2.2 kV, MS full scans were performed in the ultra-high-field Orbitrap mass analyzer in range of m/z 300-1400 with a resolution of 120,000 at m/z 200, the maximum injection time (MIT) was 80 ms and the automatic gain control (AGC) was set as 3×10^6^. The top 20 intense ions were selected and subjected to Orbitrap for further fragmentation via high energy collision dissociation (HCD) activation over a mass range between m/z 200 and 2000 at a resolution of 15,000 with the intensity threshold kept at 2.6×10^4^. We selected ions with charge state from 2+ to 6+ for sequencing. Normalized collision energy (NCE) was set at 27. For each MS_2_ scan, the target AGC was set as 2 × 10^4^ and the MIT was 19 ms. The dynamic exclusion for precursor ions were set over a time window of 12 s to suppress repeated peak fragmentation.

### Proteomics and Phosphoproteomics Data Processing

Raw data of proteomics and phosphoproteomics were processed with MaxQuant (version 1.6.0.1). MS and MS/MS spectra were searched against the human Uniprot database (version 201506) using the Andromeda search engine. The database search was performed with the following parameters: an initial mass tolerance of ±20 ppm and a final mass tolerance of ±0.5 Da for precursor masses, ±0.6 Da for HCD ion trap fragment ions, and two missed cleavages allowed. Cysteine carbamidomethylation was used as a fixed modification, and methionine oxidation, protein N-terminal acetylation, and serine, threonine, and tyrosine phosphorylation were included as variable modifications for phosphoproteomics data. The false discovery rate was set at 0.01 for peptides, proteins, and phosphosites; the minimum peptide length allowed was seven amino acids, and a minimum Andromeda peptide score of 60 was required. The match-between-runs feature was enabled. A site localization probability of at least 0.75 and a score difference of at least 5 were used as thresholds for the localization of phosphoresidues.

Reverse and contaminated proteins were removed after database search. Next, the proteins and phosphosites with quantitative intensity values in less than 4 of the 21 samples in each time point were removed. After the filtration, the quantitative protein information was median standardized. Normalization was performed by subtracting the median of log transformed intensities for each nano-LC-MS/MS run. Given that the data followed the normal distribution, we imputed the missing values from normal distribution in Perseus software ^47^. To identify significantly regulated phosphorylation sites or proteins, limma package was used for HepG2 cell with coumarin treatment. A fold change of > 1.5 or < 0.67 with p-value < 0.05 was used as the cut off threshold for differentially expressed proteins and phosphosites. As no biological replicates were included in the doxorubicin experiments on AC16 cells, we used a more stringent 2-fold change for differential expression of genes/proteins/phosphosites. The Mfuzz package was used to obtain the temporal trends of changes in phospho-sites.

KEGG enrichment analysis was analyzed and visualized in the DAVID bioinformatics database. The change at a functional level in different groups was performed with reactome analysis. Boxplot and PCA plots were generated using *ggplot2* ^48^. A pairwise Pearson correlation coefficient was calculated for all group runs in *R* v.3.6.1^49^. Different colors of sample correlation analysis were used to represent different sample types in the hierarchical clustering analysis by using the ward.2 method in the *pheatmap* package.

### Western blotting

After cell treatment with the same conditions as in the proteomics study. An equal amount of total protein was resolved by a 10% sodium dodecyl sulfate polyacrylamide gel (SDS-PAGE) and transferred to nitrocellulose membranes (Bio-Rad, Hercules, CA, USA). Membranes were blocked and incubated with either anti Phospho-(Ser/Thr) Phe Antibody (9631S) or anti Phospho-Tyrosine Rabbit pAb (AP0905) primary antibodies overnight at 4 °C, then washed and subsequently incubated with goat anti-mouse IgG-HRP (CW0102) secondary antibodies. GAPDH was used to control the amount of proteins loaded for western blotting, in which anti rabbit IgG-HRP (CW0103S) was used as the primary antibodies. The membranes were stained with SuperSignal™ West Pico PLUS Chemiluminescent Substrate (34580, Thermo Fisher Scientific, San Jose, CA, USA). Protein bands were visualized using Tanon-5200 (Tanon, Shanghai, China).

### Computational modeling

A minimal mathematical model of posttranslational and transcriptional stress adaptation was constructed to illustrate the differential responses. Model parameter values and ordinary differential equations (ODEs) are provided in Tables S15 and S16 respectively. The model was constructed and simulated in Berkeley Madonna (version 8.3.18, University of California, Berkeley, CA, USA) using the “Rosenbrock (stiff)” ODE solver. The model code was provided as supplemental material.

## Result

### Quality assurance of multi-omics study for cellular response to coumarin

After evaluating the cellular response data for coumarin in the literature^50–52^, we found that the lowest AC_50_ (concentration inducing the half-maximum assay response) for coumarin in HepG2 cells is about 30 µM (Table S1). Since it was anticipated that the proteome and phosoproteome are sensitive to chemical perturbations, we decided to use a wide range of concentrations biased to the lower end of 30 µM to explore the coumarin’s effects at these two omics levels as well as on the transcriptome. Besides the vehicle control, 6 concentrations were used in HepG2 cells ranging between 0.001-100 µM at a 10-fold increment (Figure 1). This range of coumarin concentrations in our previous study caused no detected changes in cell count and viability of HepG2 cells^39^. In addition, we defined 0 μM as control, 0.001-0.01 μM as low-concentration, 0.1-1 μM as mid-concentration, and 10-100 μM as high-concentration groups. For transcriptomics analysis, the cells were treated for 6 and 24 hours for each coumarin concentration. For proteomics and phosphoproteomics analyses, the cells were treated for 10 min and 24 hours. The 10 min time point was chosen because rapid changes in the phosoproteome, and possibly the proteome, are expected to occur on this time scale^53^.

**Figure 1.**
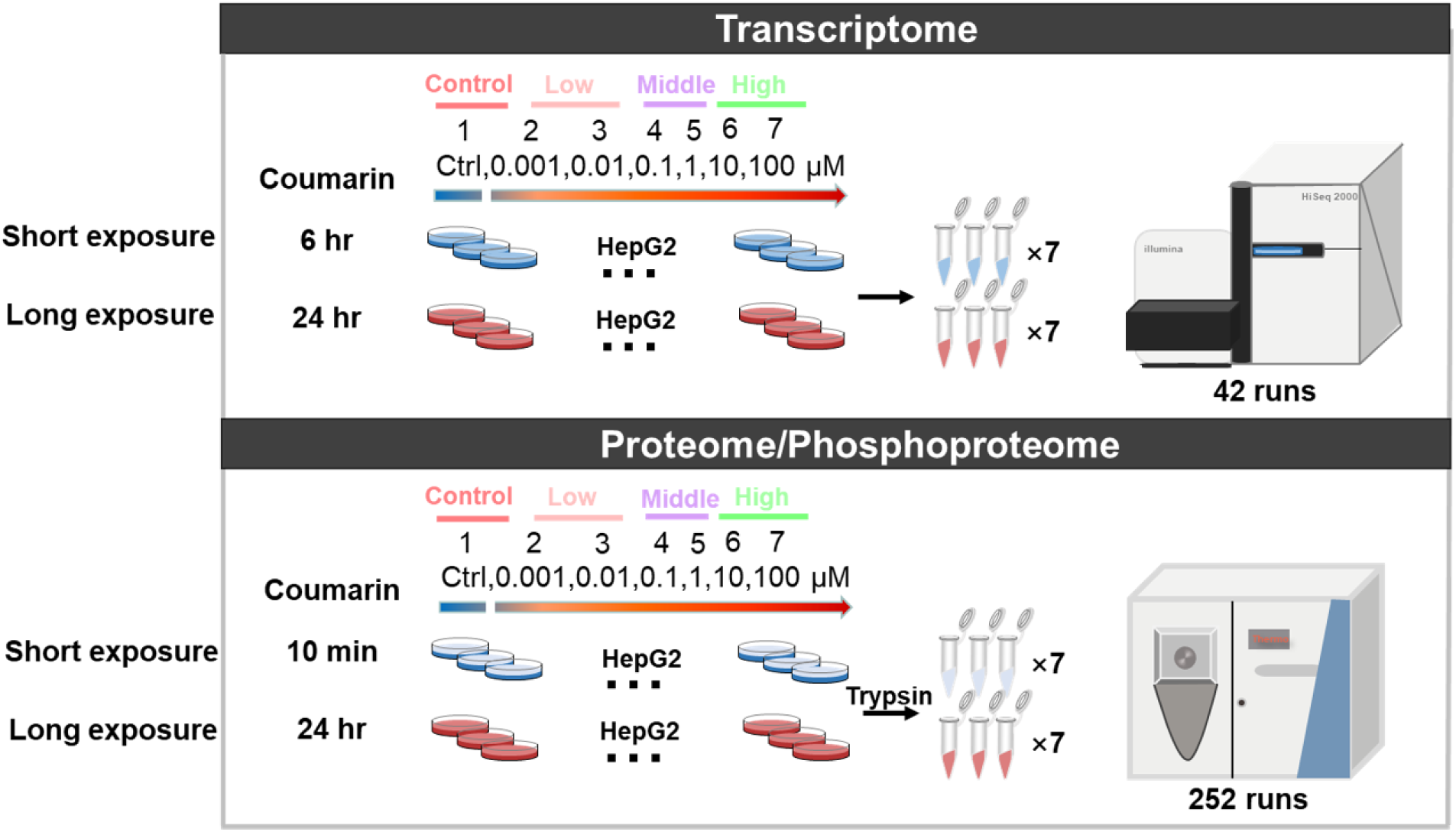
The experimental procedures for multi-omics measurements of cellular response to coumarin. The workflow of omics study for cellular response of coumarin treated HepG2 cells. In the transcriptomics study, in addition to vehicle control, HepG2 cells were treated with 6 concentrations of coumarin ranging from 0.001 to 100 μM as indicated. The concentrations are numerically coded as 1-7 as indicated for ease of referencing in other figures. The cells were recovered for RNAseq after treated for 6 and 24 hours, which were denoted as short-exposure and long-exposure, respectively. Each concentration-time experimental condition was repeated for three biological replicates. A total of 42 samples were collected for RNA-seq. In proteomics and phosphoproteomics studies all experimental conditions remained the same except that the treatment durations are 10 min and 6 hours.

### Global differences in transcriptomics, proteomics and phosphoproteomics responses to coumarin

After the strict quality control on multi-omics data were conducted (Figures S1-4, Tables S2-6), we performed PCA analysis which was considered as an initial effort to investigate which omics is better at distinguishing different levels of perturbations by coumarin, we first applied principal component analysis (PCA) to the datasets. For the 6 hours exposure groups in the RNAseq experiment, PCA can barely separate the 21 samples treated with different concentrations of coumarin, except that the three replicates treated with the highest concentration, i.e., 100 µM, tended to cluster together and stayed away from the remaining samples (Figure 2A). The segregation of the 100-µM samples from the rest is even more obvious in the 24 hours exposure group (Figure 2B). These results are consistent with the reported AC50 values for coumarin in HepG2 cells, which ranged between 28-163 µM, indicating that transcriptomics changes are likely concordant with those cellular assay changes.

**Figure 2.**
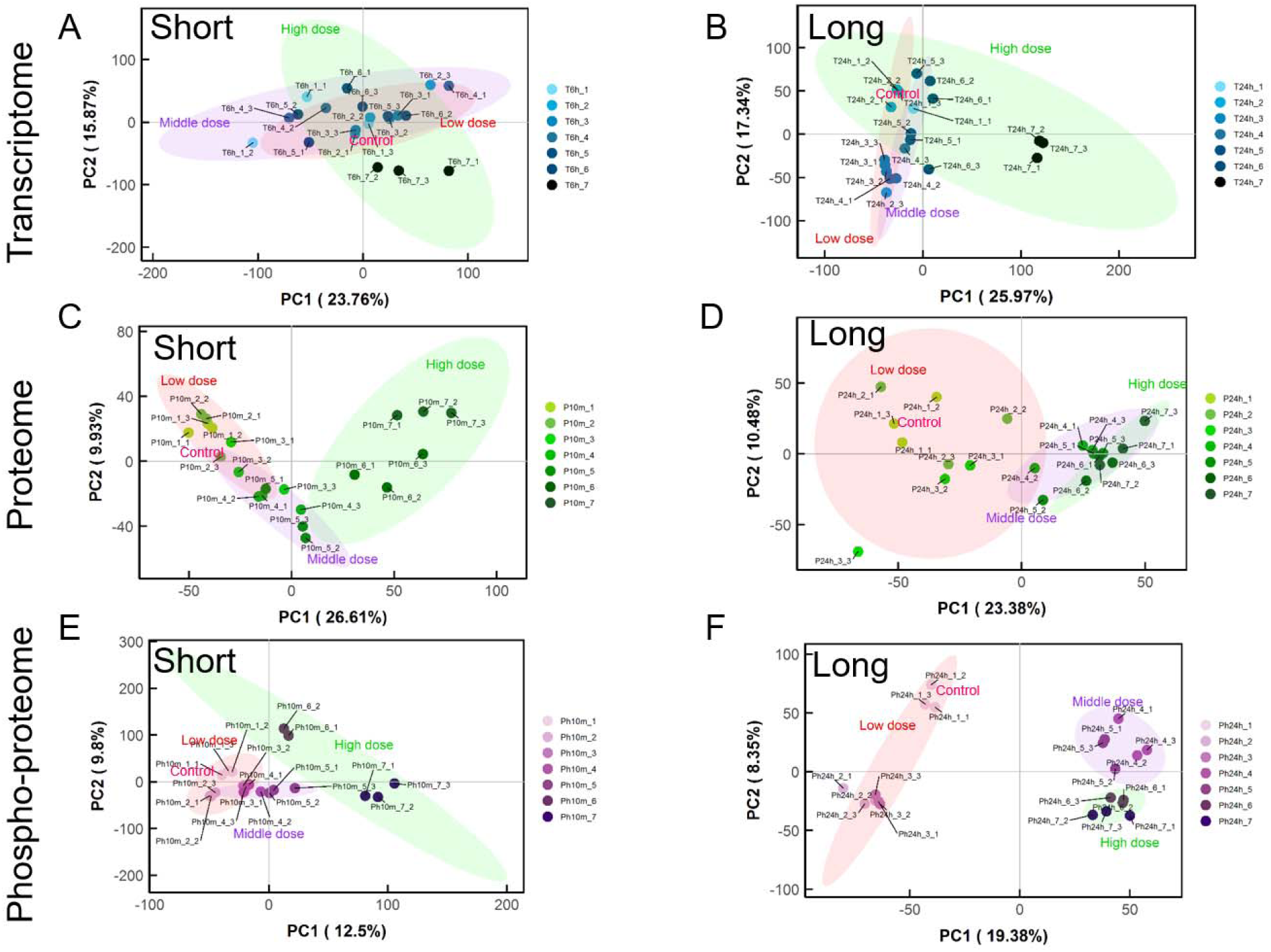
PCA analysis of omics data generated from HepG2 cells treated with various concentrations of coumarin. (A & B) PCA analysis of transcriptomics data for short (6-hr) and long (24-hr) exposure to coumarin, respectively. (C & D) PCA analysis of proteomics data for short (10-min) and long (24-hr) exposure to coumarin, respectively. (E & F) PCA analysis of phosphoproteomics data for short (10 min) and long (24 hours) exposure to coumarin, respectively. Color-shaded ellipses encircle different coumarin dose groups: red, control group and low-dose (0-0.01 µM); pink, middle-dose (0.1-1 µM); and green, high-dose (10-100 µM).

In contrast to the results obtained with transcriptomics, the proteome is better at distinguishing different coumarin concentrations, especially for high and low concentrations. Even for 10 min exposure, both the 10 and 100 μM samples are easily separated from the remaining concentrations and the three 10 μM replicates form a cluster that can be distinguished from the 100 μM replicate cluster (Figure 2C). There also seems to be a better separation between the middle-concentration (0.1-1 μM) and low-concentration samples (0.001-0.01 μM) of the proteomics data than the transcriptomics data. Interestingly, for 24 hours exposure, the high-concentration samples can no longer be easily differentiated from the middle-concentration samples. However, the low-concentration group can be readily distinguished from middle and high-concentration groups (Figure 2D). In addition, it is interesting to found that the results of control samples were quite close to the low concentration samples which not only because of the limit of detection sensitivity for mass spectrum, but may also indicate the biological significances between them and turn out to be consistent with our knowledge that the biological activity of low-dose of coumarin were slight^54^. Similar to the proteomics results, the phosphoproteome could effectively distinguish high and low-concentration groups. For 10 min exposure to coumarin, the high-concentration group stands out from the rest, and the low and middle-concentration groups also seem distinguishable from each other (Figure 2E). 24 hours exposure gave the clearest distinction, where all three dose groups are clearly separated on the 2D PC plot, and the samples treated with 0.001-0.01 μM coumarin are also distanced from the vehicle control (Ph24h_1_x) samples (Figure 2F). In summary, the proteome and phosphoproteome are much more sensitive than the transcriptome in distinguishing cellular responses to coumarin exposure of different concentrations.

To further compare the effects of treatments with different concentrations of coumarin for different exposure durations, we applied unsupervised hierarchical clustering to the dataset. For short exposure duration, as expected, the transcriptomics data could not distinguish coumarin concentration groups as they all are mixed together (Figure S5A). With the long exposure duration, only the 100 μM coumarin group could be separated into a distinguishable cluster from the rest of the concentration groups (Figure S5B). The proteome data could divide into two broad clusters, one for high-concentration coumarin treatment, another for mid and low-concentration coumarin treatments and control group, and the latter can be further grouped into separate clusters (Figure S5C). The control and the ultra-low (0.001 μM) samples cannot be readily distinguished from each other. With the exposure prolonged to 24 hours, different coumarin concentration groups can still be distinguished but the separation does not seem to improve over the short exposure data (Figure S5D). Similar to the PCA results, the phosphoproteome showed the most distinct dose dependent separation among the three omics datasets. The replicates for each coumarin concentration are always cleanly clustered together, for both the short and long exposure durations. The separation and similarity between treatment groups strictly follow the order of the coumarin concentrations including the control (Figure S5E&F). Taken together, the hierarchical clustering results are consistent with the results from PCA analysis, further confirming that the phosphoproteome could effectively distinguish the cellular responses to chemical treatments at different concentrations, especially for ultra-low concentrations.

### Phosphoproteome is several orders of magnitude more sensitive than transcriptome in response to coumarin

To further understand the sensitivity of various omics techniques in detecting chemical perturbations, volcano plots were generated to identify significantly altered mRNA, protein, and phosphosite features in HepG2 cells (Figure S6). The log_2_ fold change of each feature in a coumarin concentration-duration treatment group relative to control was plotted against –log_10_ of the p-value for the transcriptomics, proteomics, and phosphoproteomics data, as shown in Figure S6A-6C, respectively. As reflected by the widening of the volcano plots, a striking trend is that the number of significantly altered features increases tremendously moving from transcriptomics, to proteomics and phosphoproteomics responses (Figure 3A). As the coumarin concentration increases, the numbers of significantly altered features increase, which also seem to increase with longer exposure duration, especially for the mRNA data. When the numbers of significantly altered features achieved by different coumarin concentrations were normalized to that achieved at 100 μM, the highest coumarin concentration, each omics exhibits distinct concentration-response profiles (Figure 3B&C). The transcriptome is clearly the least sensitive, where a high response only takes off at >10 μM. In contrast, the phosphoproteomics response is the most sensitive, already at 50% or higher at 0.001 μM coumarin for both short and long exposure durations. Compared with transcriptomics and phosphoproteomics responses, the sensitivity of the proteomics response falls in between.

**Figure 3.**
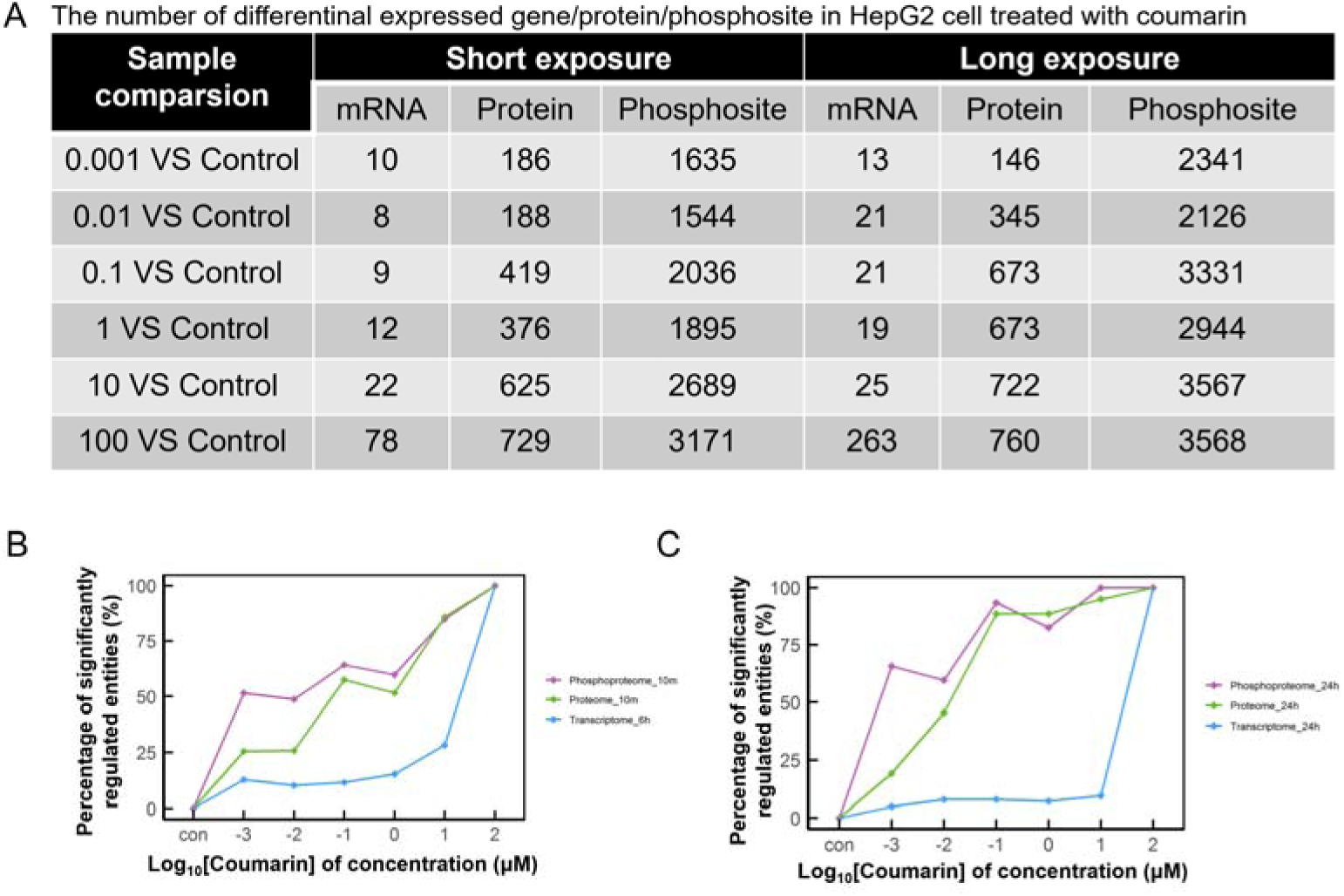
Phosphoproteome is the most sensitive omics marker to detect cellular responses to low concentrations of coumarin in HepG2 cells. (A) The number of differentially altered features relative to control in the transcriptome, proteome and phosphoproteome of HepG2 cells treated with coumarin of various concentrations for short (10-30 min) or long durations (24 hours). (B & C) Percentage of differentially altered features normalized to the number of differentially altered features at the highest concentration (100 μM) for short (B) and long (C) exposures, respectively.

### High detection sensitivity is conserved for phosphoproteomics in doxorubicin-treated AC16 cells

We next investigated whether the high sensitivity of phosphoproteomics observed for cellular response to coumarin in HepG2 cells can also be recapitulated in other cell types in response to other chemicals. We applied the same experimental approach to AC16 cells, a cardiomyocyte cell line, treated with doxorubicin. Doxorubicin is a chemotherapeutic agent used in the treatment of solid tumors and hematological malignancies. The most important dose-limiting toxicity of doxorubicin is its cardiotoxicity^55^. The previously reported PoD of doxorubicin in AC16 cells was 125 nM, at which a variety of cellular responses were significantly altered, including mitochondrial ROS, membrane potential and DNA content, and key genes involved in mitochondrial biogenesis and antioxidant response^56^. Similarly, iPSC-derived cardiomyocytes experienced beating changes at around 150 nM of doxorubicin treatment, which is a PoD also eliciting transcriptomics changes^57^. To test the distinguishing power of different omics in detecting perturbation by doxorubicin, we selected 10 nM as the concentration to treat AC16 cells, which is at least 10-folder lower than the PoD above. The omic-level responses were measured at varied exposure durations, including 10, 30 and 360 min, which covered both fast and delayed responses (Figure 4A).

**Figure 4.**
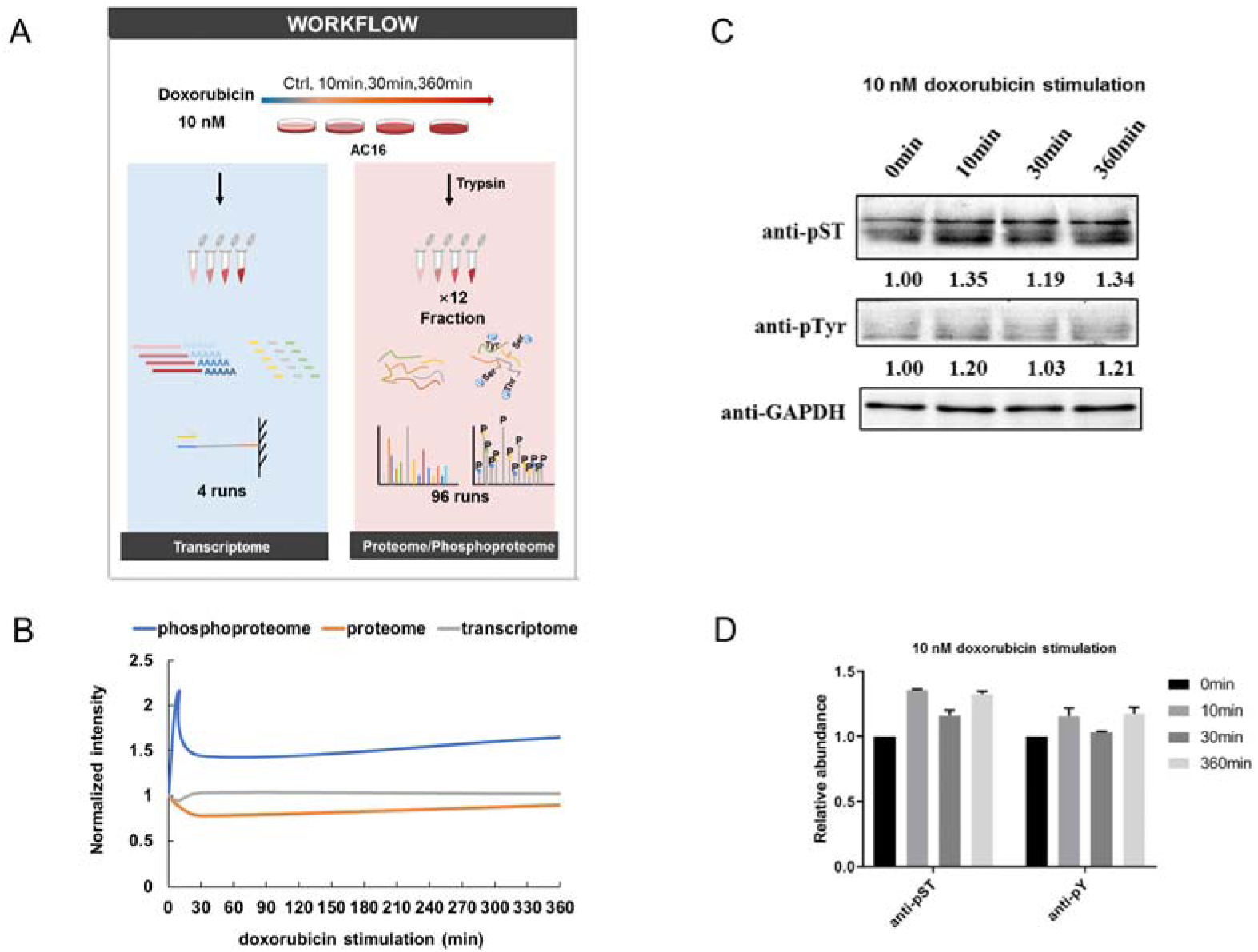
Phosphoproteome is the most sensitive omics marker to detect cellular responses to doxorubicin at a low concentration in AC16 cells. (A) The workflow of omics study for cellular response of AC16 cells to 10 nM doxorubicin treatment for 0, 10, 30 and 360 min. A total of 4 samples were collected for RNA sequencing. For proteomics and phosphoproteomics, each of the sample had 12 fractions. (B) The relative intensity of abundance normalized to control at 0 min for transcriptome, proteome and phosphoproteome. (C) Western blot analysis of phospho-protein in total cell lysates extracted from AC16 cells treated with 10 nM doxorubicin for 10, 30, and 360 min. Anti-pST and anti-pTyr: pan antibodies used to detect phosphorylated serine/threonine and tyrosine respectively. (D) Quantification of (C).

The qualities of the transcriptomics, proteomics and phosphoproteomics data of doxorubicin-treated AC16 cells were very similar to those of coumarin-treated HepG2 samples (Figure S7-S9, Table S7-S9). Given the high quality and the consistency of the replicates of the coumarin-treated HepG2 samples, we are confident with the low variability of our data-generating platform and moved forward with single replicates for the doxorubicin-treated AC16 cells. After normalizing the abundance, the global signals for transcriptomics and proteomics at various time points were stable compared with the control. This result is consistent with the fact that the concentration of doxorubicin used here is much lower than the PoD concentration and thus is not expected to elicit responses at transcriptomics and proteomics levels. In contrast, in response to 10 nM doxorubicin, the abundance signal of altered phosphoproteome spiked immediately to a peak level at 10 min, then decreased till 30 min, and increased slightly by 360 min (Figure 4B). Probing total cell lysate with pan-antibodies against pSer/pThr (pST) and pTyr, we found that anti-pST signal was increased in doxorubicin treated samples compared with the control, but not pTyr (Figure 4C&D). This might result from low detection sensitivity of pan-antibodies against pTyr used in this study^58, 59^. Taken together, these results confirmed that phosphoproteome was more sensitive to reflect cellular response under low and ultra-low concentrations of chemicals that normally do not disturb the transcriptome and proteome.

### Potential sensitive biomarkers for cellular response to doxorubicin

Finally, we asked the question whether we could find specific sensitive biomarkers in the phosphoproteome as a mechanistic indicator for doxorubicin. We systematically analyzed the multi-omics data generated from the doxorubicin-treated AC16 samples. By using a stringent two-fold change as the threshold, we identified 695, 729 and 686 significantly up-regulated phosphosites for samples treated with doxorubicin for 10, 30, and 360 min, respectively. The down-regulated phosphosites were 629, 640 and 742, respectively (Figure 5A). Reactome pathway analysis^60^ of all up-regulated phosphosites showed that the cell cycle and mitosis related pathways were significantly enriched (underlined in Figure 5B). Other altered pathways include those for RNA metabolism and Rho GTPase signaling. This is consistent with the observations that doxorubicin inhibited cell proliferation through cell cycle arrest at the G2/M phase^61^ and doxorubicin induced increases in mitotic activity in BE(2)-C neuroblastoma^62^. However, this cell cycle effect occurs at much lower concentrations than therapeutic concentrations.

**Figure 5.**
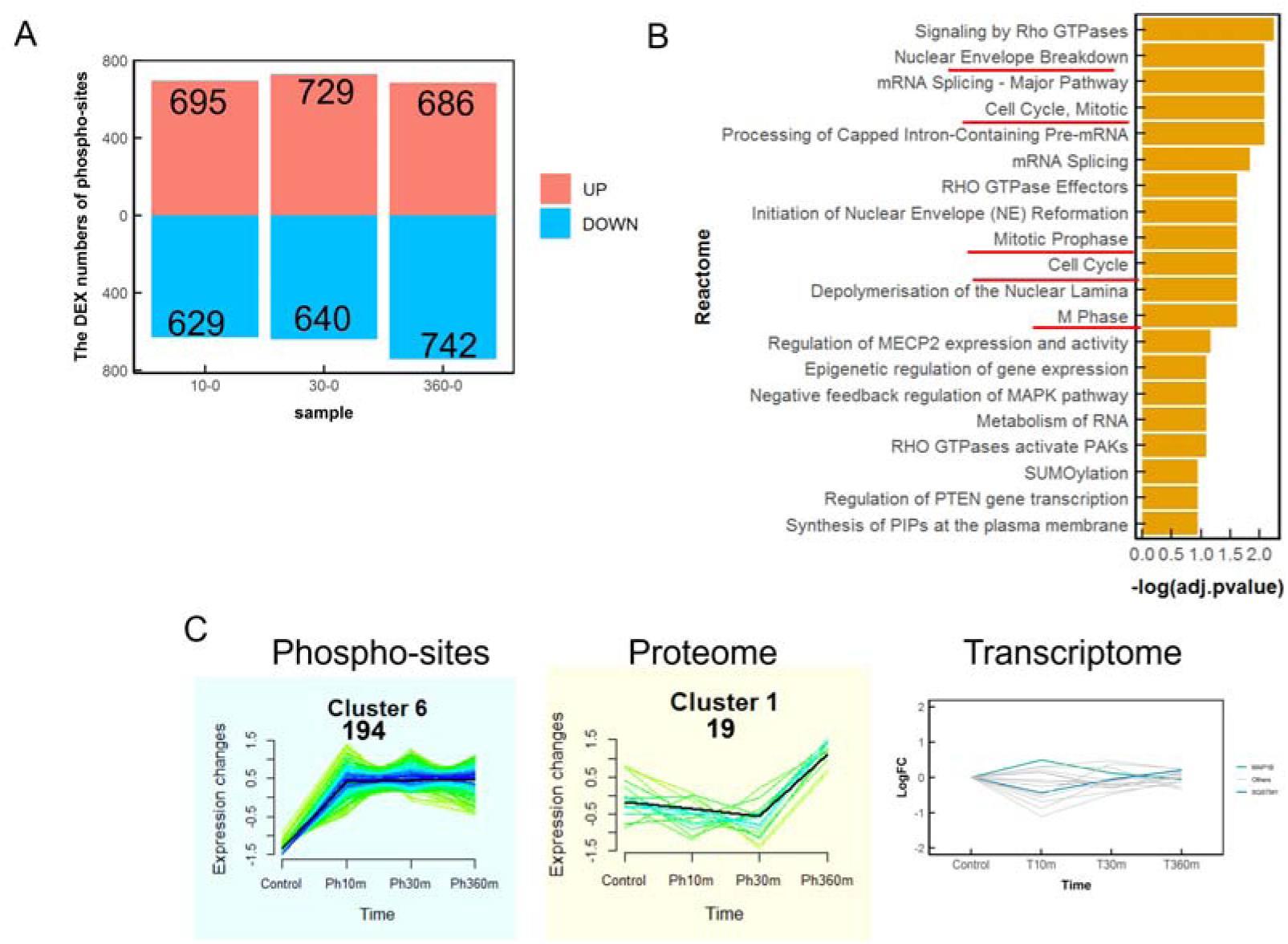
Phosphoproteomics revealed sensitive pathway enrichment in low-concentration doxorubicin-treated AC16 cells. (A) The number of significantly up- and down-regulated phosphosites in AC16 cells treated with 10 nM doxorubicin for 10, 30, and 360 min (with FC > 2 or < 0.5). (B) GSEA-based enrichment analysis of GO biological processes up-regulated by10 nM doxorubicin treatment. All of the up-regulated phosphoproteins were combined for this analysis and top 20 Reactome pathways were displayed. (C) A total of 194 phosphosites were upregulated persistently throughout the 360-min duration of doxorubicin treatment. 19 proteins identified in the proteomics study form a cluster that exhibited upregulation only at 360 min. No significant changes in transcriptome were identified among the corresponding genes.

Time-series clustering analysis showed that a total of 194 phosphosites were upregulated persistently throughout the 360-min duration of doxorubicin treatment (Figure 5C, Table S10). These upregulated phosphosites correspond to 167 proteins, among which, 157 were also identified in the proteome dataset (Table S11). Interestingly, 19 of the proteins form a single proteomics cluster, which exhibited baseline-level responses in the first 30 min and then upregulation at 6 hours (Figure 5C). In contrast, no tangible changes in the corresponding genes were observed through the entire time course (Figure 5C). These results suggest that upregulation of these genes only occurred at the posttranscriptional, and especially posttranslational, level.

We next examined how some well-known doxorubicin-induced pathways behave in the three omics datasets. By eliciting oxidative stress, doxorubicin can induce the antioxidant response in cardiomyocytes, including the master transcription factor Nrf2 and signature target gene HMOX1^8^. Although Nrf2 is activated through protein stabilization and in some cases transcriptionally through autoregulation^63, 64^, it was not identified in either of our omics datasets. It is because the concentration of doxorubicin used is well below the PoD. In comparison, HMOX1, which is a highly sensitive biomarker for antioxidant response, was identified in our proteome data (Table S12). TSC2 (ser939) is an important protein that can activate p53, which mediates the DNA damage response to doxorubicin^65^. TSC2 was identified in our phosphoproteomics data (Table S13). All of these three known molecular markers have been identified and included in these 19 proteins. These results strongly supported that phosphoproteomics is a sensitive approach to assess the cellular response to doxorubicin. STQSTM1/p62 is heavily related to cardiotoxicity although without any direct evidence regarding the toxicity of doxorubicin so far^66, 67^. The phosphorylation of p62 is controlled by CDK1 and plays biological roles in promoting cell cycle initiation, mitosis and tumor proliferation^68^. p62 (Ser28) was increased about 19-fold at 10 min compared with control, which is consistent with the global trend of doxorubicin-induced phosphoproteomics changes (Table S14). The biological function of this molecule in the cellular response to doxorubicin treatment is unknown.

### A minimal mathematical model recapitulated tiered cross-omics responses

To better interpret and understand the observed temporal and concentration-dependent multi-omics response patterns, we constructed a minimal mathematical model capturing the framework of tiered cross-omics cellular adaptation to stress^5, 32^. The model contains both post translationally and transcriptionally-mediated negative feedbacks (Figure 6A, Table S15-16). Specifically, it includes phosphorylation and thus posttranslational activation of a preexisting, basal stress protein by a kinase which is activated by controlled state (such as ROS and DNA damage) when perturbed by stressor (step 1 in Figure 6A), transcriptional induction of the mRNA and protein of the stress gene by a transcription factor that is activated by the altered controlled state (step 2), and lastly a global translational inhibition of proteins nonessential to the stress response but necessary for cell functions (step 3). The model recapitulated the general pattern of the dynamic and concentration-response of the transcriptome, proteome and phosphoproteome observed in the present study. The transient spikes of the active phospho-stress protein (Ga) at early timepoints which return to lower steady-state levels upon mild stress (Figure 6B) are consistent with the temporal profile of the phosphoproteome observed in doxorubicin-treated HepG2 cells (Figure 4B). Such dynamics are due to the initial perturbation by the stressor S and subsequent engagement of the posttranslational feedback that brings the controlled state (Y) to a nearly basal level. During this time frame of 6 hours, no transcription of mRNA (M) is tangibly induced. Simulation of 24 hours exposure to the stressor resulted in concentration-response of different sensitivity (Figure 6C). The active stress protein (Ga) exhibited a response that arises at a low S level, while the M and total protein responses takes off at a much higher S level around 10. At this S level, the cell function protein (CFP) expression begins to be repressed with the controlled state Y deviating from the baseline (Figure 6D), demarcating a PoD that may be linked to apical endpoint changes.

**Figure 6.**
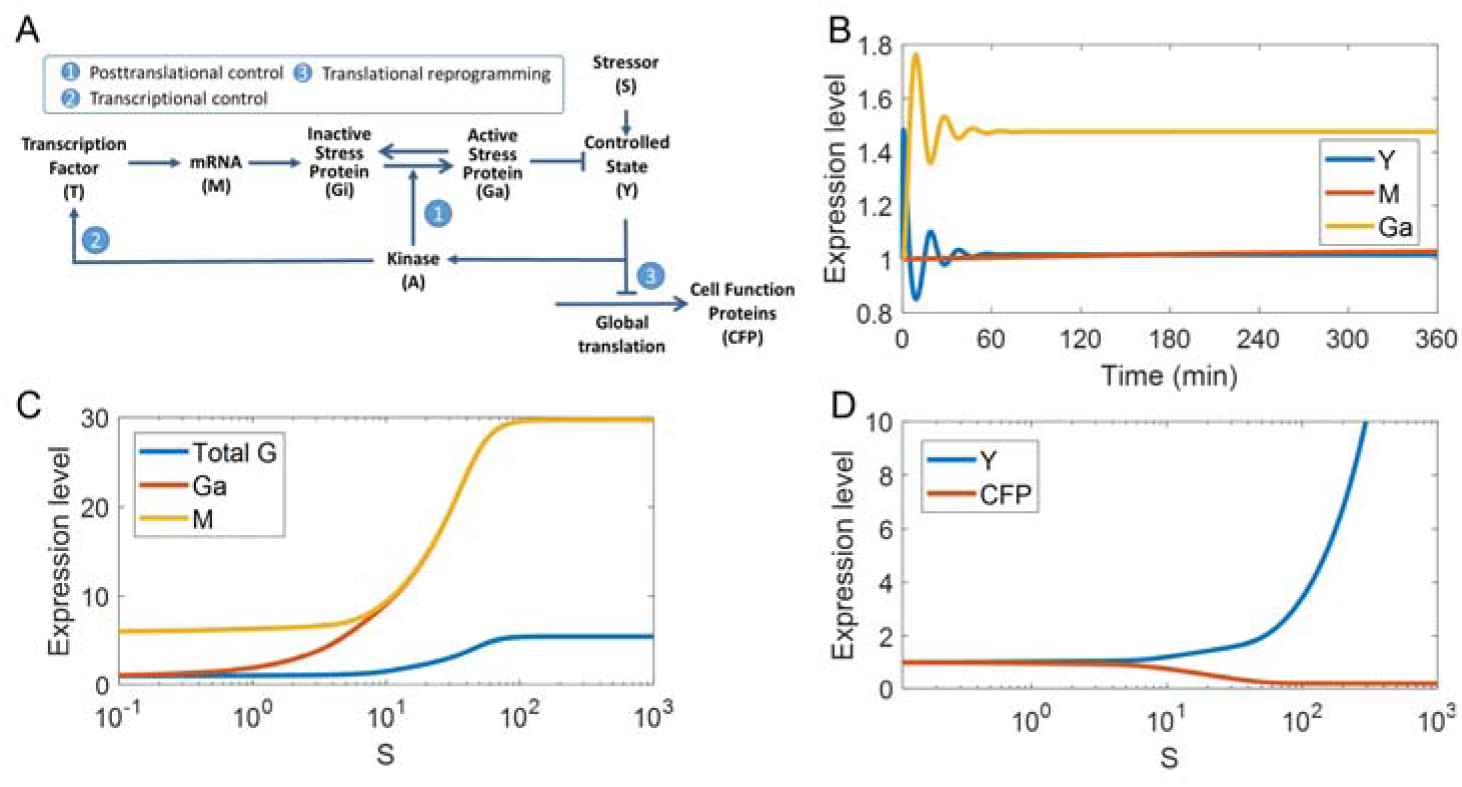
Simulations of cell stress response by two-tiered transcriptional and posttranslational control. (A) Mode of cellular stress response to stressor S may involve both transcriptional and posttranslational control of the cellular state (Y). The transcriptional induction of mRNA (M) for stress gene (G) is activated by transcription factor (T) through pathway #2. Posttranslational modification control bypasses the slow-acting transcriptional loop by regulating the activities of preexisting inactive stress proteins (Gi) through phosphorylation to convert Gi into active stress protein (Ga) via pathway #1. The stress can also directly affect specialized cell function proteins (CFP) through step #3. Pointed arrows denote activation and blunted arrows denote inhibition. (B) Temporal responses of Y, Ga and M in response to a mild S level (0.5). (C) Dose responses of M, Ga and total G after 24-h exposure. (D) Dose responses of Y and CFP after 24-h exposure.

## Discussion

The transcriptome, proteome, and phosphoproteome of the cell have unique properties, with information constantly flowing across these scales of biology. In the present study, we demonstrated with coumarin and doxorubicin, two compounds with fundamentally distinct molecular initiating events and modes of action, that phosphoproteomics changes occur not only earlier in time as expected, but also at lower chemical concentrations, compared with proteomics and transcriptomics alterations. As a result, changes in the phosphoproteome can provide sensitive markers compared with changes in the proteome and transcriptome to indicate alterations in cellular state following chemical exposure. Although only two compounds were examined here, to our knowledge, this is the first study that compared multi-omics changes across many concentrations and time points with replicates.

A single-omics technique such as in vitro high throughput transcriptomics has recently been shown to be suitable for NAM-based hazard evaluation of environmental chemicals^69^. In particular, high throughput transcriptomics has been used in lower tiers of NGRA assessments as a conservative estimate of a quantitative POD in the ‘protection not prediction paradigm’ with the aim of casting the broadest net possible for capturing the potential relevant bioactivity associated with chemical treatment^70^. However, the use of a single omics techniques is unlikely to lead to systematic understanding of pathways’ response to chemical exposure and hence elucidate on the mode of action of the substance. This is because a single-omics technique measures biomolecule of a specific type, e.g. RNAs in case of transcriptomics, and thus captures time-specific changes only for a small subset of components of a particular pathway. Therefore, where residual uncertainty in a safety assessment means that such mechanistic understanding is needed, other omics technologies may provide valuable data.

Proteomics techniques have developed rapidly over the past decade and are now being able to identify and quantify several thousands of proteins from a single sample. Proteins represent the biomolecules that directly operate in the cells and are therefore closer to the functional level. Several studies have shown transcript expression levels for a specific transcript product may not correlate to its protein expression level^71–74^. Indeed, besides the transcriptionally controlled effects on protein abundance, there are translational and posttranslational effects that are not reflected on the transcriptome level. There are many different types of PTMs but the most abundant PTM is phosphorylation that has been the subject of this work. Sampadi et al has recently demonstrated in a phosphoproteomics study that chemical stressors with different modes of action can induce distinctive and complex phosphorylation signaling patterns^75^. In this work we were able to detect another layer of biological response upon chemical exposure, by identifying differentially expressed proteins and phosphosites.

In the era of NAM, a WoE approach that integrates multiple data streams is increasingly adopted in the Next Generation Risk Assessement. Transcriptomics has gained tremendous traction as an emerging high-coverage omics platform^76^ and its importance to mechanistically-based safety assessment should be better appreciated and interpreted. Confidence in the use of transcriptomically-determined PoDs will increase if mechanistic justifications can be provided in the larger context of cellular responses to chemical perturbations. This is of particular importance in cases when NAMs used in lower tiers of risk assessment do not provide sufficient information to make a decision on the absence of biological activity for a given exposure scenario. In these cases, while transcriptionally-mediated cytoprotective gene induction is well understood, other protective mechanisms that operate before transcriptional induction is initiated are no less important biologically and can be exploited as biomarkers to assess the degree of cellular perturbation. Here we compared changes across transcriptomic, proteomic and phosphoproteomic measurements obtained with many replicates, concentration and time points to understand the sensitivity of the three omics techniques in response to a toxicant. We have demonstrated that phosphoproteomics changes occur not only earlier in time as expected but also at low chemical concentrations and hence are proximal to the very early effects induced by chemical treatment. Therefore, phosphoproteomics and proteomics changes may play an essential role in investigating mechanisms of early adaptation and protection in cells. Such non-transcriptomic omics data can be cross-referenced with and corroborate the transcriptomically defined PoD as part of the multi-omics WoE approach.

Although far from being as sensitive as the phosphoproteome, the proteome exhibited abundance changes of hundreds of proteins as early as 10 min in HepG2 cells after coumarin treatment (Figure 3). Protein half-lives in mammalian cells are generally in the order of hours on average, therefore at 10 min of exposure, little change would be expected in protein abundance. However, proteins can differ in turnover rates dramatically, across several orders of magnitude^77^, and many of them are shorted-lived^53^. Protein stability is also highly regulated, many with half-lives shorter than 20 min or even as short as a few minutes^78–81^. Therefore, if coumarin treatment leads to altered protein synthesis or degradation, protein abundance changes can be observed in 10 min for fast-turnover proteins.

The current analysis showed that integration of different omics layers to be used in toxicology is not straightforward and indeed poses many challenges as recently reviewed^82^. One of the challenges represents the choice of time points for different experiments. The choice of time points used in this study has been based on the previous reports^8, 9^, although we are aware that this is a complex issue and there is not only a time variance between different omics layers, e.g. proteome is downstream of the transcriptome, but also different biological processes occur on different time scale within a single omics layer, e.g. translation occurs on a different time scale from cell division. In addition, since proteomics and phosphoproteomics have not been routinely used as NAM-based tool in hazard evaluation, their reproducibility of data acquisition and data analysis demand more consideration. Furthermore, the coverage of expressed proteins, phosphosites and transcripts needs to be addressed in order to ensure that relevant biology has been sufficiently covered. Finally, the application of such a multi-omics approach to case studies plays a critical role in building capability, understanding and ultimately confidence in bioactivity characterization as integrating multi-omics is beginning to be considered in toxicity testing and drug development^11, 12, 82, 83^. Detailed pathway analyses of the phosphoproteomics, proteomics, and transcriptomics dataset reported in this study and derivation of associated PoDs at different omics levels will be reported in separate upcoming publications to further reveal mechanistic underpinnings for the tiered adaptive cellular response.

In summary, we propose to further evaluate the potential of multi-omics and its application in Next Generation Risk Assessment in future using case study chemicals. We have presented promising results by assessing proteomics and phosphoproteomics along with transcriptomics and strongly believe that such integration of multiple omics data sets will offer a substantial improvement in detecting molecular responses due to more complete representation of the perturbed toxicity pathways.

## Acknowledgments

We are indebted to Doctors Chao Liu, Liang Zhao, Yanan Yin, Xuehui Peng for support in the early stage of this project. We thank Doctors Yingchun Wang, Yangjun Zhang and Guibin Wang on the help of reagents, peptide manual checking and LC-MS analysis of proteomics and phosphoproteomics samples. This study was funded by the Unilever (MA-2018−02170N), MOST (2017YFA0505100 and 2020YFE0202200), the National Natural Science Foundation of China (NSFC) (32141003, 32070668, 31870824, 91839302, 31901037, and 32071431), the Innovation Foundation of Medicine (AWS17J008, 19SWAQ17 and 20SWAQX34), and the Foundation of State Key Lab of Proteomics (SKLP-K201704 and SKLP-K201901), the CAMS Innovation Fund for Medical Sciences (2019-I2M-5-017), the Beijing-Tianjin-Hebei Basic Research Cooperation Project (J200001), the Mass Spectrometry Platform Open Project of National Center for Protein Sciences Beijing and Unilever-Emory Collaborative Project (MA-2015-02026).

## Supplemental Figures

Figure S1. Quality control for the transcriptomics study in HepG2 cells treated with Coumarin.

Figure S2. Quality control for the proteomics study in HepG2 cells treated with Coumarin.

Figure S3. Quality control for the phospho-proteomics study in HepG2 cells treated with Coumarin.

Figure S4. Pair-wise Pearson correlation analysis of phosphoproteomics, proteomics, and transcriptomics samples of HepG2 cells treated with Coumarin.

Figure S5. Hierarchical clustering of the transcriptome, proteome, and phosphoproteome of HepG2 cells treated with Coumarin.

Figure S6. Volcano plots of differentially expressed features in the proteome, phosphoproteome, and transcriptome of HepG2 cells treated with Coumarin.

Figure S7. Quality control for the transcriptomics study in AC16 cells treated with 10 nM doxorubicin for 10, 30, and 360 min.

Figure S8. Quality control for the proteomics study in AC16 cells treated with 10 nM doxorubicin for 10, 30, and 360 min.

Figure S9. Quality control for the phosphor-proteomics study in AC16 cells treated with 10 nM doxorubicin for 10, 30, and 360 min.

## Supplemental Tables

Table S1. The AC50 values of coumarin on HepG2 cells in references

Table S2. The genes identified from HepG2 cells treated with different concentration of coumarin.

Table S3. The proteins identified from HepG2 cells treated with different concentration of coumarin for 10 min.

Table S4. The proteins identified from HepG2 cells treated with different concentration of coumarin for 24 hours.

Table S5. The phosphosites identified from HepG2 cells treated with different concentrations of coumarin for 10 min.

Table S6. The phosphosites identified from HepG2 cells treated with different concentration of coumarin for 24 hours.

Table S7. The genes identified from AC16 cells treated with doxorubicin for different times.

Table S8. The proteins identified from AC16 cells treated with doxorubicin for different times

Table S9. The phosphosites identified from AC16 cells treated with doxorubicin for different times.

Table S10. The number of 194 phosphosites (cluster 6) identified from AC16 cells treated with doxorubicin for different times.

Table S11. The number of 157 proteins (match with 194 phosphosites) identified from AC16 cells treated with doxorubicin for different times.

Table S12. The HMOX1 proteins identified from AC16 cells treated with doxorubicin for different times.

Table S13. The phosphodites of TSC2 identified from AC16 cells treated with doxorubicin for different times.

Table S14. The phosphodites of SQSTM1 identified from AC16 cells treated with doxorubicin for different times.

Table S15. Parameter values of the mathematical model.

Table S16. Ordinary differential equations (ODEs) of the mathematical model.

## References

(1) Paul Friedman, K.; Gagne, M.; Loo, L.-H.; Karamertzanis, P.; Netzeva, T.; Sobanski, T.; Franzosa, J. A.; Richard, A. M.; Lougee, R. R.; Gissi, A.; Lee, J.-Y. J.; Angrish, M.; Dorne, J. L.; Foster, S.; Raffaele, K.; Bahadori, T.; Gwinn, M. R.; Lambert, J.; Whelan, M.; Rasenberg, M.; Barton-Maclaren, T.; Thomas, R. S. Utility of in vitro bioactivity as a lower bound estimate of in vivo adverse effect levels and in risk-based prioritization. Toxicological Sciences. 2019, 173, (1), 202–225.

(2) Krewski, D.; Andersen, M. E.; Tyshenko, M. G.; Krishnan, K.; Hartung, T.; Boekelheide, K.; Wambaugh, J. F.; Jones, D.; Whelan, M.; Thomas, R.; Yauk, C.; Barton-Maclaren, T.; Cote, I. Toxicity testing in the 21st century: Progress in the past decade and future perspectives. Archives of Toxicology. 2020, 94, (1), 1–58.

(3) Dent, M. P.; Vaillancourt, E.; Thomas, R. S.; Carmichael, P. L.; Ouedraogo, G.; Kojima, H.; Barroso, J.; Ansell, J.; Barton-Maclaren, T. S.; Bennekou, S. H.; Boekelheide, K.; Ezendam, J.; Field, J.; Fitzpatrick, S.; Hatao, M.; Kreiling, R.; Lorencini, M.; Mahony, C.; Montemayor, B.; Mazaro-Costa, R.; Oliveira, J.; Rogiers, V.; Smegal, D.; Taalman, R.; Tokura, Y.; Verma, R.; Willett, C.; Yang, C. Paving the way for application of next generation risk assessment to safety decision-making for cosmetic ingredients. Regulatory Toxicology and Pharmacology. 2021, 125, 105026.

(4) Simmons, S. O.; Fan, C. Y.; Ramabhadran, R. Cellular stress response pathway system as a sentinel ensemble in toxicological screening. Toxicol Sci. 2009, 111, (2), 202–25.

(5) Zhang, Q.; Bhattacharya, S.; Pi, J.; Clewell, R. A.; Carmichael, P. L.; Andersen, M. E. Adaptive posttranslational control in cellular stress response pathways and its relationship to toxicity testing and safety assessment. Toxicol Sci. 2015, 147, (2), 302–16.

(6) Bergamini, G.; Bell, K.; Shimamura, S.; Werner, T.; Cansfield, A.; Müller, K.; Perrin, J.; Rau, C.; Ellard, K.; Hopf, C.; Doce, C.; Leggate, D.; Mangano, R.; Mathieson, T.; O’Mahony, A.; Plavec, I.; Rharbaoui, F.; Reinhard, F.; Savitski, M. M.; Ramsden, N.; Hirsch, E.; Drewes, G.; Rausch, O.; Bantscheff, M.; Neubauer, G. A selective inhibitor reveals pi3kγ dependence of t(h)17 cell differentiation. Nat Chem Biol. 2012, 8, (6), 576–82.

(7) Hendriks, G.; Derr, R. S.; Misovic, B.; Morolli, B.; Calléja, F. M.; Vrieling, H. The extended toxtracker assay discriminates between induction of DNA damage, oxidative stress, and protein misfolding. Toxicol Sci. 2016, 150, (1), 190–203.

(8) Yuan, H.; Zhang, Q.; Guo, J.; Zhang, T.; Zhao, J.; Li, J.; White, A.; Carmichael, P. L.; Westmoreland, C.; Peng, S. A pgc-1α-mediated transcriptional network maintains mitochondrial redox and bioenergetic homeostasis against doxorubicin-induced toxicity in human cardiomyocytes: Implementation of tt21c. Toxicol Sci. 2016, 150, (2), 400–17.

(9) Baltazar, M. T.; Cable, S.; Carmichael, P. L.; Cubberley, R.; Cull, T.; Delagrange, M.; Dent, M. P.; Hatherell, S.; Houghton, J.; Kukic, P.; Li, H.; Lee, M. Y.; Malcomber, S.; Middleton, A. M.; Moxon, T. E.; Nathanail, A. V.; Nicol, B.; Pendlington, R.; Reynolds, G.; Reynolds, J.; White, A.; Westmoreland, C. A next-generation risk assessment case study for coumarin in cosmetic products. Toxicol. Sci. 2020, 176, (1), 236–252.

(10) Thomas, R. S.; Bahadori, T.; Buckley, T. J.; Cowden, J.; Deisenroth, C.; Dionisio, K. L.; Frithsen, J. B.; Grulke, C. M.; Gwinn, M. R.; Harrill, J. A.; Higuchi, M.; Houck, K. A.; Hughes, M. F.; Hunter, E. S.; Isaacs, K. K.; Judson, R. S.; Knudsen, T. B.; Lambert, J. C.; Linnenbrink, M.; Martin, T. M.; Newton, S. R.; Padilla, S.; Patlewicz, G.; Paul-Friedman, K.; Phillips, K. A.; Richard, A. M.; Sams, R.; Shafer, T. J.; Setzer, R. W.; Shah, I.; Simmons, J. E.; Simmons, S. O.; Singh, A.; Sobus, J. R.; Strynar, M.; Swank, A.; Tornero-Valez, R.; Ulrich, E. M.; Villeneuve, D. L.; Wambaugh, J. F.; Wetmore, B. A.; Williams, A. J. The next generation blueprint of computational toxicology at the u.S. Environmental protection agency. Toxicol Sci. 2019, 169, (2), 317–332.

(11) Wilmes, A.; Bielow, C.; Ranninger, C.; Bellwon, P.; Aschauer, L.; Limonciel, A.; Chassaigne, H.; Kristl, T.; Aiche, S.; Huber, C. G.; Guillou, C.; Hewitt, P.; Leonard, M. O.; Dekant, W.; Bois, F.; Jennings, P. Mechanism of cisplatin proximal tubule toxicity revealed by integrating transcriptomics, proteomics, metabolomics and biokinetics. Toxicol In Vitro. 2015, 30, (1 Pt A), 117–27.

(12) Wilmes, A.; Limonciel, A.; Aschauer, L.; Moenks, K.; Bielow, C.; Leonard, M. O.; Hamon, J.; Carpi, D.; Ruzek, S.; Handler, A.; Schmal, O.; Herrgen, K.; Bellwon, P.; Burek, C.; Truisi, G. L.; Hewitt, P.; Di Consiglio, E.; Testai, E.; Blaauboer, B. J.; Guillou, C.; Huber, C. G.; Lukas, A.; Pfaller, W.; Mueller, S. O.; Bois, F. Y.; Dekant, W.; Jennings, P. Application of integrated transcriptomic, proteomic and metabolomic profiling for the delineation of mechanisms of drug induced cell stress. J Proteomics. 2013, 79, 180–94.

(13) Hudson, K. M.; Shiver, E.; Yu, J.; Mehta, S.; Jima, D. D.; Kane, M. A.; Patisaul, H. B.; Cowley, M. Transcriptomic, proteomic, and metabolomic analyses identify candidate pathways linking maternal cadmium exposure to altered neurodevelopment and behavior. Scientific Reports. 2021, 11, (1), 16302.

(14) Gatzidou, E. T.; Zira, A. N.; Theocharis, S. E. Toxicogenomics: A pivotal piece in the puzzle of toxicological research. J Appl Toxicol. 2007, 27, (4), 302–9.

(15) Van Hummelen, P.; Sasaki, J. State-of-the-art genomics approaches in toxicology. Mutat Res. 2010, 705, (3), 165–71.

(16) Chepelev, N. L.; Moffat, I. D.; Labib, S.; Bourdon-Lacombe, J.; Kuo, B.; Buick, J. K.; Lemieux, F.; Malik, A. I.; Halappanavar, S.; Williams, A.; Yauk, C. L. Integrating toxicogenomics into human health risk assessment: Lessons learned from the benzo[a]pyrene case study. Crit. Rev. Toxicol. 2015, 45, (1), 44–52.

(17) Farmahin, R.; Williams, A.; Kuo, B.; Chepelev, N. L.; Thomas, R. S.; Barton-Maclaren, T. S.; Curran, I. H.; Nong, A.; Wade, M. G.; Yauk, C. L. Recommended approaches in the application of toxicogenomics to derive points of departure for chemical risk assessment. Arch Toxicol. 2017, 91, (5), 2045–2065.

(18) Thomas, R. S.; Wesselkamper, S. C.; Wang, N. C.; Zhao, Q. J.; Petersen, D. D.; Lambert, J. C.; Cote, I.; Yang, L.; Healy, E.; Black, M. B.; Clewell, H. J., 3rd; Allen, B. C.; Andersen, M. E. Temporal concordance between apical and transcriptional points of departure for chemical risk assessment. Toxicol. Sci. 2013, 134, (1), 180–94.

(19) Moffat, I.; Chepelev, N.; Labib, S.; Bourdon-Lacombe, J.; Kuo, B.; Buick, J. K.; Lemieux, F.; Williams, A.; Halappanavar, S.; Malik, A.; Luijten, M.; Aubrecht, J.; Hyduke, D. R.; Fornace, A. J., Jr.; Swartz, C. D.; Recio, L.; Yauk, C. L. Comparison of toxicogenomics and traditional approaches to inform mode of action and points of departure in human health risk assessment of benzo[a]pyrene in drinking water. Crit. Rev. Toxicol. 2015, 45, (1), 1–43.

(20) Zhou, Y. H.; Cichocki, J. A.; Soldatow, V. Y.; Scholl, E. H.; Gallins, P. J.; Jima, D.; Yoo, H. S.; Chiu, W. A.; Wright, F. A.; Rusyn, I. Comparative dose-response analysis of liver and kidney transcriptomic effects of trichloroethylene and tetrachloroethylene in b6c3f1 mouse. Toxicol Sci. 2017, 160, (1), 95–110.

(21) Dean, J. L.; Zhao, Q. J.; Lambert, J. C.; Hawkins, B. S.; Thomas, R. S.; Wesselkamper, S. C. Application of gene set enrichment analysis for identification of chemically induced, biologically relevant transcriptomic networks and potential utilization in human health risk assessment. Toxicol Sci. 2017, 157, (1), 85–99.

(22) Bhat, V. S.; Hester, S. D.; Nesnow, S.; Eastmond, D. A. Concordance of transcriptional and apical benchmark dose levels for conazole-induced liver effects in mice. Toxicol Sci. 2013, 136, (1), 205–15.

(23) Harrill, J. A.; Everett, L. J.; Haggard, D. E.; Sheffield, T.; Bundy, J. L.; Willis, C. M.; Thomas, R. S.; Shah, I.; Judson, R. S. High-throughput transcriptomics platform for screening environmental chemicals. Toxicological Sciences. 2021, 181, (1), 68–89.

(24) Yang, L.; Allen, B. C.; Thomas, R. S. Bmdexpress: A software tool for the benchmark dose analyses of genomic data. BMC genomics. 2007, 8, (1), 1–8.

(25) Phillips, J. R.; Svoboda, D. L.; Tandon, A.; Patel, S.; Sedykh, A.; Mav, D.; Kuo, B.; Yauk, C. L.; Yang, L.; Thomas, R. S.; Gift, J. S.; Davis, J. A.; Olszyk, L.; Merrick, B. A.; Paules, R. S.; Parham, F.; Saddler, T.; Shah, R. R.; Auerbach, S. S. Bmdexpress 2: Enhanced transcriptomic dose-response analysis workflow. Bioinformatics. 2019, 35, (10), 1780–1782.

(26) Buttgereit, F.; Brand, M. D. A hierarchy of atp-consuming processes in mammalian cells. Biochem J. 1995, 312 *( Pt* *1*), 163–7.

(27) Hoffmann, F.; Rinas, U. On-line estimation of the metabolic burden resulting from the synthesis of plasmid-encoded and heat-shock proteins by monitoring respiratory energy generation. Biotechnol. Bioeng. 2001, 76, (4), 333–40.

(28) Klaassen, C. D.; Liu, J. Induction of metallothionein as an adaptive mechanism affecting the magnitude and progression of toxicological injury. Environ Health Perspect. 1998, 106 *Suppl 1*, 297–300.

(29) Pi, J.; Zhang, Q.; Woods, C. G.; Wong, V.; Collins, S.; Andersen, M. E. Activation of nrf2-mediated oxidative stress response in macrophages by hypochlorous acid. Toxicol Appl Pharmacol. 2008, 226, (3), 236–43.

(30) Sokolova, I. M. Energy-limited tolerance to stress as a conceptual framework to integrate the effects of multiple stressors. Integr Comp Biol. 2013, 53, (4), 597–608.

(31) Chadwick, J. G., Jr.; Nislow, K. H.; McCormick, S. D. Thermal onset of cellular and endocrine stress responses correspond to ecological limits in brook trout, an iconic cold-water fish. Conserv Physiol. 2015, 3, (1), cov017.

(32) Zhang, Q.; Li, J.; Middleton, A.; Bhattacharya, S.; Conolly, R. B. Bridging the data gap from in vitro toxicity testing to chemical safety assessment through computational modeling. Frontiers in Public Health. 2018, 6, (261).

(33) Mendes, C.; Serpa, J. Metabolic remodelling: An accomplice for new therapeutic strategies to fight lung cancer. Antioxidants (Basel). 2019, 8, (12).

(34) Rein, T. Post-translational modifications and stress adaptation: The paradigm of fkbp51. Biochem Soc Trans. 2020, 48, (2), 441–449.

(35) Gitan, R. S.; Eide, D. J. Zinc-regulated ubiquitin conjugation signals endocytosis of the yeast zrt1 zinc transporter. Biochem J. 2000, 346 *Pt* *2*, 329–36.

(36) Dihazi, H.; Kessler, R.; Eschrich, K. High osmolarity glycerol (hog) pathway-induced phosphorylation and activation of 6-phosphofructo-2-kinase are essential for glycerol accumulation and yeast cell proliferation under hyperosmotic stress. J. Biol. Chem. 2004, 279, (23), 23961–8.

(37) Krejsa, C. M.; Franklin, C. C.; White, C. C.; Ledbetter, J. A.; Schieven, G. L.; Kavanagh, T. J. Rapid activation of glutamate cysteine ligase following oxidative stress. J. Biol. Chem. 2010, 285, (21), 16116–24.

(38) Zhao, M.; Wu, F.; Xu, P. Development of a rapid high-efficiency scalable process for acetylated sus scrofa cationic trypsin production from escherichia coli inclusion bodies. Protein expression and purification. 2015, 116.

(39) Hatherell, S.; Baltazar, M. T.; Reynolds, J.; Carmichael, P. L.; Dent, M.; Li, H.; Ryder, S.; White, A.; Walker, P.; Middleton, A. M. Identifying and characterizing stress pathways of concern for consumer safety in next-generation risk assessment. Toxicological Sciences. 2020, 176, (1), 11–33.

(40) Li, Y.; Dammer, E. B.; Gao, Y.; Lan, Q.; Villamil, M. A.; Duong, D. M.; Zhang, C.; Ping, L.; Lauinger, L.; Flick, K.; Xu, Z.; Wei, W.; Xing, X.; Chang, L.; Jin, J.; Hong, X.; Zhu, Y.; Wu, J.; Deng, Z.; He, F.; Kaiser, P.; Xu, P. Proteomics links ubiquitin chain topology change to transcription factor activation. Mol. Cell. 2019, 76, (1), 126–137.e7.

(41) Pertea, M.; Kim, D.; Pertea, G. M.; Leek, J. T.; Salzberg, S. L. Transcript-level expression analysis of rna-seq experiments with hisat, stringtie and ballgown. Nature protocols. 2016, 11, (9), 1650–67.

(42) Xu, P.; Duong, D. M.; Peng, J. Systematical optimization of reverse-phase chromatography for shotgun proteomics. Journal of proteome research. 2009, 8, (8), 3944–50.

(43) Wu, F.; Zhao, M.; Zhang, Y.; Su, N.; Xiong, Z.; Xu, P. Recombinant acetylated trypsin demonstrates superior stability and higher activity than commercial products in quantitative proteomics studies. Rapid communications in mass spectrometry : RCM. 2016, 30, (8), 1059–66.

(44) Peng, X.; Xu, F.; Liu, S.; Li, S.; Huang, Q.; Chang, L.; Wang, L.; Ma, X.; He, F.; Xu, P. Identification of missing proteins in the phosphoproteome of kidney cancer. Journal of proteome research. 2017, 16, (12), 4364–4373.

(45) Xu, F.; Yu, L.; Peng, X.; Zhang, J.; Li, S.; Liu, S.; Yin, Y.; An, Z.; Wang, F.; Fu, Y.; Xu, P. Unambiguous phosphosite localization through the combination of trypsin and lysarginase mirror spectra in a large-scale phosphoproteome study. Journal of proteome research. 2020, 19, (6), 2185–2194.

(46) Zhai, L.; Chang, C.; Li, N.; Duong, D. M.; Chen, H.; Deng, Z.; Yang, J.; Hong, X.; Zhu, Y.; Xu, P. Systematic research on the pretreatment of peptides for quantitative proteomics using a cJJ microcolumn. Proteomics. 2013, 13, (15), 2229–37.

(47) Tyanova, S.; Temu, T.; Sinitcyn, P.; Carlson, A.; Hein, M. Y.; Geiger, T.; Mann, M.; Cox, J. The perseus computational platform for comprehensive analysis of (prote)omics data. Nature methods. 2016.

(48) Wickham, H. *Ggplot2: Elegant graphics for data analysis*. Springer Publishing Company, Incorporated: 2009.

(49) Team, R. C. R: A language and environment for statistical computing. 2013.

(50) Romanov, S.; Medvedev, A.; Gambarian, M.; Poltoratskaya, N.; Moeser, M.; Medvedeva, L.; Gambarian, M.; Diatchenko, L.; Makarov, S. Homogeneous reporter system enables quantitative functional assessment of multiple transcription factors. Nature methods. 2008, 5, (3), 253–60.

(51) Martin, M. T.; Dix, D. J.; Judson, R. S.; Kavlock, R. J.; Reif, D. M.; Richard, A. M.; Rotroff, D. M.; Romanov, S.; Medvedev, A.; Poltoratskaya, N.; Gambarian, M.; Moeser, M.; Makarov, S. S.; Houck, K. A. Impact of environmental chemicals on key transcription regulators and correlation to toxicity end points within epa’s toxcast program. Chemical Research in Toxicology. 2010, 23, (3), 578–590.

(52) Giuliano, K. A.; Gough, A. H.; Taylor, D. L.; Vernetti, L. A.; Johnston, P. A. Early safety assessment using cellular systems biology yields insights into mechanisms of action. Journal of biomolecular screening. 2010, 15, (7), 783–97.

(53) Li, J.; Cai, Z.; Vaites, L. P.; Shen, N.; Mitchell, D. C.; Huttlin, E. L.; Paulo, J. A.; Harry, B. L.; Gygi, S. P. Proteome-wide mapping of short-lived proteins in human cells. Molecular Cell. 2021, 81, (22), 4722–4735.e5.

(54) Hatherell, S.; Baltazar, M. T.; Reynolds, J.; Carmichael, P. L.; Dent, M.; Li, H.; Ryder, S.; White, A.; Walker, P.; Middleton, A. M. Identifying and characterizing stress pathways of concern for consumer safety in next-generation risk assessment. Toxicol. Sci. 2020, 176, (1), 11–33.

(55) Lenaz, L.; Page, J. A. Cardiotoxicity of adriamycin and related anthracyclines. Cancer Treatment Reviews. 1976, 3, (3), 111–120.

(56) Yuan, H.; Zhang, Q.; Guo, J.; Zhang, T.; Zhao, J.; Li, J.; White, A.; Carmichael, P. L.; Westmoreland, C.; Peng, S. A pgc-1α-mediated transcriptional network maintains mitochondrial redox and bioenergetic homeostasis against doxorubicin-induced toxicity in human cardiomyocytes: Implementation of tt21c. Toxicological Sciences. 2016, 150, (2), 400–417.

(57) Chaudhari, U.; Nemade, H.; Wagh, V.; Gaspar, J. A.; Ellis, J. K.; Srinivasan, S. P.; Spitkovski, D.; Nguemo, F.; Louisse, J.; Bremer, S.; Hescheler, J.; Keun, H. C.; Hengstler, J. G.; Sachinidis, A. Identification of genomic biomarkers for anthracycline-induced cardiotoxicity in human ipsc-derived cardiomyocytes: An in vitro repeated exposure toxicity approach for safety assessment. Archives of Toxicology. 2016, 90, (11), 2763–2777.

(58) Bergström Lind, S.; Molin, M.; Savitski, M. M.; Emilsson, L.; Aström, J.; Hedberg, L.; Adams, C.; Nielsen, M. L.; Engström, A.; Elfineh, L.; Andersson, E.; Zubarev, R. A.; Pettersson, U. Immunoaffinity enrichments followed by mass spectrometric detection for studying global protein tyrosine phosphorylation. J. Proteome Res. 2008, 7, (7), 2897–910.

(59) Bian, Y.; Li, L.; Dong, M.; Liu, X.; Kaneko, T.; Cheng, K.; Liu, H.; Voss, C.; Cao, X.; Wang, Y.; Litchfield, D.; Ye, M.; Li, S. S.; Zou, H. Ultra-deep tyrosine phosphoproteomics enabled by a phosphotyrosine superbinder. Nat. Chem. Biol. 2016, 12, (11), 959–966.

(60) Jassal, B.; Matthews, L.; Viteri, G.; Gong, C.; Lorente, P.; Fabregat, A.; Sidiropoulos, K.; Cook, J.; Gillespie, M.; Haw, R.; Loney, F.; May, B.; Milacic, M.; Rothfels, K.; Sevilla, C.; Shamovsky, V.; Shorser, S.; Varusai, T.; Weiser, J.; Wu, G.; Stein, L.; Hermjakob, H.; D’Eustachio, P. The reactome pathway knowledgebase. Nucleic Acids Res. 2020, 48, (D1), D498–D503.

(61) Kim, H. S.; Lee, Y. S.; Kim, D. K. Doxorubicin exerts cytotoxic effects through cell cycle arrest and fas-mediated cell death. Pharmacology. 2009, 84, (5), 300–309.

(62) Hultman, I.; Haeggblom, L.; Rognmo, I.; Jansson Edqvist, J.; Blomberg, E.; Ali, R.; Phillips, L.; Sandstedt, B.; Kogner, P.; Shirazi Fard, S.; Ährlund-Richter, L. Doxorubicin-provoked increase of mitotic activity and concomitant drain of g0-pool in therapy-resistant be(2)-c neuroblastoma. PLOS ONE. 2018, 13, (1), e0190970.

(63) Kobayashi, A.; Kang, M. I.; Watai, Y.; Tong, K. I.; Shibata, T.; Uchida, K.; Yamamoto, M. Oxidative and electrophilic stresses activate nrf2 through inhibition of ubiquitination activity of keap1. Molecular and cellular biology. 2006, 26, (1), 221–9.

(64) Kwak, M. K.; Itoh, K.; Yamamoto, M.; Kensler, T. W. Enhanced expression of the transcription factor nrf2 by cancer chemopreventive agents: Role of antioxidant response element-like sequences in the nrf2 promoter. Molecular and cellular biology. 2002, 22, (9), 2883–92.

(65) Feng, Z.; Zhang, H.; Levine, A. J.; Jin, S. The coordinate regulation of the p53 and mtor pathways in cells. Proceedings of the National Academy of Sciences of the United States of America. 2005, 102, (23), 8204–8209.

(66) Li, D. L.; Wang, Z. V.; Ding, G.; Tan, W.; Luo, X.; Criollo, A.; Xie, M.; Jiang, N.; May, H.; Kyrychenko, V.; Schneider, J. W.; Gillette, T. G.; Hill, J. A. Doxorubicin blocks cardiomyocyte autophagic flux by inhibiting lysosome acidification. Circulation. 2016, 133, (17), 1668–87.

(67) Li, R.; Huang, Y.; Semple, I.; Kim, M.; Zhang, Z.; Lee, J. H. Cardioprotective roles of sestrin 1 and sestrin 2 against doxorubicin cardiotoxicity. Am J Physiol Heart Circ Physiol. 2019, 317, (1), H39–H48.

(68) Linares, J. F.; Amanchy, R.; Greis, K.; Diaz-Meco, M. T.; Moscat, J. Phosphorylation of p62 by cdk1 controls the timely transit of cells through mitosis and tumor cell proliferation. Molecular and cellular biology. 2011, 31, (1), 105–17.

(69) Auerbach, S. S.; Paules, R. S. Genomic dose response: Successes, challenges, and next steps. Current Opinion in Toxicology. 2018, 11-12, 84–92.

(70) Harrill, J.; Shah, I.; Setzer, R. W.; Haggard, D.; Auerbach, S.; Judson, R.; Thomas, R. S. Considerations for strategic use of high-throughput transcriptomics chemical screening data in regulatory decisions. Curr Opin Toxicol. 2019, 15, 64–75.

(71) Grün, D.; Kirchner, M.; Thierfelder, N.; Stoeckius, M.; Selbach, M.; Rajewsky, N. Conservation of mrna and protein expression during development of c. elegans. Cell Reports. 2014, 6, (3), 565-577.

(72) Maier, T.; Güell, M.; Serrano, L. Correlation of mrna and protein in complex biological samples. FEBS Lett. 2009, 583, (24), 3966–73.

(73) Gygi, S. P.; Rochon, Y.; Franza, B. R.; Aebersold, R. Correlation between protein and mrna abundance in yeast. Mol Cell Biol. 1999, 19, (3), 1720–30.

(74) Futcher, B.; Latter, G. I.; Monardo, P.; McLaughlin, C. S.; Garrels, J. I. A sampling of the yeast proteome. Mol Cell Biol. 1999, 19, (11), 7357–68.

(75) Sampadi, B.; Pines, A.; Munk, S.; Mišovic, B.; de Groot, A. J.; van de Water, B.; Olsen, J. V.; Mullenders, L. H. F.; Vrieling, H. Quantitative phosphoproteomics to unravel the cellular response to chemical stressors with different modes of action. Arch. Toxicol. 2020, 94, (5), 1655–1671.

(76) Harrill, J. A.; Viant, M. R.; Yauk, C. L.; Sachana, M.; Gant, T. W.; Auerbach, S. S.; Beger, R. D.; Bouhifd, M.; O’Brien, J.; Burgoon, L.; Caiment, F.; Carpi, D.; Chen, T.; Chorley, B. N.; Colbourne, J.; Corvi, R.; Debrauwer, L.; O’Donovan, C.; Ebbels, T. M. D.; Ekman, D. R.; Faulhammer, F.; Gribaldo, L.; Hilton, G. M.; Jones, S. P.; Kende, A.; Lawson, T. N.; Leite, S. B.; Leonards, P. E. G.; Luijten, M.; Martin, A.; Moussa, L.; Rudaz, S.; Schmitz, O.; Sobanski, T.; Strauss, V.; Vaccari, M.; Vijay, V.; Weber, R. J. M.; Williams, A. J.; Williams, A.; Thomas, R. S.; Whelan, M. Progress towards an oecd reporting framework for transcriptomics and metabolomics in regulatory toxicology. Regulatory Toxicology and Pharmacology. 2021, 125, 105020.

(77) Schwanhäusser, B.; Busse, D.; Li, N.; Dittmar, G.; Schuchhardt, J.; Wolf, J.; Chen, W.; Selbach, M. Corrigendum: Global quantification of mammalian gene expression control. Nature. 2013, 495, (7439), 126–7.

(78) Berra, E.; Roux, D.; Richard, D. E.; Pouysségur, J. Hypoxia-inducible factor-1 alpha (hif-1 alpha) escapes o(2)-driven proteasomal degradation irrespective of its subcellular localization: Nucleus or cytoplasm. EMBO Rep. 2001, 2, (7), 615–20.

(79) Khalil, H. S.; Goltsov, A.; Langdon, S. P.; Harrison, D. J.; Bown, J.; Deeni, Y. Quantitative analysis of nrf2 pathway reveals key elements of the regulatory circuits underlying antioxidant response and proliferation of ovarian cancer cells. J Biotechnol. 2015, 202, 12–30.

(80) Heinzel, S.; Binh Giang, T.; Kan, A.; Marchingo, J. M.; Lye, B. K.; Corcoran, L. M.; Hodgkin, P. D. A myc-dependent division timer complements a cell-death timer to regulate t cell and b cell responses. Nat Immunol. 2017, 18, (1), 96–103.

(81) Levine, A. J. Targeting therapies for the p53 protein in cancer treatments. Annual Review of Cancer Biology. 2019, 3, (1), 21–34.

(82) Canzler, S.; Schor, J.; Busch, W.; Schubert, K.; Rolle-Kampczyk, U. E.; Seitz, H.; Kamp, H.; von Bergen, M.; Buesen, R.; Hackermüller, J. Prospects and challenges of multi-omics data integration in toxicology. Arch Toxicol. 2020, 94, (2), 371–388.

(83) Selevsek, N.; Caiment, F.; Nudischer, R.; Gmuender, H.; Agarkova, I.; Atkinson, F. L.; Bachmann, I.; Baier, V.; Barel, G.; Bauer, C.; Boerno, S.; Bosc, N.; Clayton, O.; Cordes, H.; Deeb, S.; Gotta, S.; Guye, P.; Hersey, A.; Hunter, F. M. I.; Kunz, L.; Lewalle, A.; Lienhard, M.; Merken, J.; Minguet, J.; Oliveira, B.; Pluess, C.; Sarkans, U.; Schrooders, Y.; Schuchhardt, J.; Smit, I.; Thiel, C.; Timmermann, B.; Verheijen, M.; Wittenberger, T.; Wolski, W.; Zerck, A.; Heymans, S.; Kuepfer, L.; Roth, A.; Schlapbach, R.; Niederer, S.; Herwig, R.; Kleinjans, J. Network integration and modelling of dynamic drug responses at multi-omics levels. Communications Biology. 2020, 3, (1), 573.

