## Supplemental materials for "The phosphoproteome is a first responder in tiered cellular adaptation to chemical stress followed by proteomics and transcriptomics alteration"

**The phosphoproteome is a first responder in tiered cellular adaption to chemical stress followed by proteomic and transcriptomic alteration**

### Supplemental methods

#### Quality control of transcriptomics, proteomics, phosphoproteomics data

For the transcriptomics study, an average of 12.3 GB RNAseq data was collected for each sample. Nearly 99% of the RNA-seq reads were mapped to the genome in all samples (Figure S1A). The number of mapped reads per sample is 79,244,722 ± 10,812,837 (mean ± SD), which are essentially constant across all sequenced samples (Figure S1B). After filtering for FPKM values greater than a stringent cutoff of 1.0 in at least one out of all 42 samples, 10,462 protein-coding genes were identified (Figure S1C, Table S2). The FPKM distributions are consistent across all samples (Figure S1D).

For the proteomics study, the generated raw MS/MS data were processed against a reference human proteome downloaded from the human Uniprot database (version 201506 ) by maxquant [55]. From the trypsin/lys-C-digested HepG2 cells, a total of 99,288 peptides were identified, which were unambiguously mapped to 6,768 proteins for the 10 min samples (Table S3), and 95,751 peptides identified and mapped to 6,537 proteins for the 24 h samples (Table S4). The mass error of most identified matched spectrum was less than 1 ppm (Figure S2A), indicating good condition of the mass spectrometer platform used. More than 5000 proteins were identified in each sample (Figure S2B). The distribution of the numbers of spectra per peptide has a mean value of 38 and the number of peptides per protein was 14, indicating the reliability of proteomic identification at the protein level (Figure S2C). The $R^2$ (Pearson) correlation value of protein abundance (LFQ) between two control replicates is 0.98 (Figure S2D), confirming the stability of our proteomics platform. After filtering and imputation, we also found that the protein abundance distributions were globally consistent in all samples (Figure S2E).

To sensitively and consistently identify phosphorylation events in HepG2 cells with or without coumarin treatment, we applied a peptide pre-separation strategy followed by enrichment of phospho-peptides in each fraction with $Ti^{4+}$-immobilized metal ion affinity chromatography (IMAC) method [44]. The generated raw MS/MS data were processed with database search. From the trypsin/lys-C-digested HepG2 cells, a total of 28,288 phosphosites and 12,799 phosphopeptides were identified, which were unambiguously mapped to 4,434 proteins for the 10 min samples (Table S5), and 24,152 phosphosites and 10,000 phosphopeptides identified and mapped to 4,155 proteins for 24 h samples (Table S6). The mass error of the most identified phosphopeptides is between -6 to 6 ppm (Figure S3A), suggesting good condition of our mass spectrometer platform. The phosphosite scores of 66.1% phosphopeptides are greater than 0.75, indicating high confidence in phosphosite localization (Figure S3B). The identified pSer, pThr and pTyr accounted for 75%, 21% and 4% of all phosphosites, respectively (Figure S3C). Among them, 62% are monophosphorylated (Figure S3D). After filtering and imputation, we also found that the distributions of the abundance of phosphosites were consistent across all samples (Figure S3E).

To further reveal the stability of the omics data across the transcriptome, proteome and phosphoproteome, we applied Pearson correlation analysis to samples of same coumarin exposure duration. The correlations are the best for the transcriptomic data, with the coefficients greater than 0.99 and 0.98 for short and long exposures, respectively (Figures S4A & S4B). The correlation coefficients for the proteomic data are greater than 0.95 and 0.93 for short and long exposure, respectively (Figure S4A & B). These results clearly illustrated that our transcriptomic and proteomic analytics was stable, which would be amenable to dose response analysis. For the phosphoproteomic data, the correlation coefficients between biological replicates receiving the same coumarin treatment in concentration and duration are greater than 0.7, which also strongly supported the stability of our proteomics platform for phosphoproteomics. However, the correlation coefficients between samples from different coumarin treatment groups are only larger than 0.5 (Figure S4A & B), which is obviously lower than those for the transcriptomics and proteomics, likely due to enrichment and higher dynamic range.

### Supplemental Figures

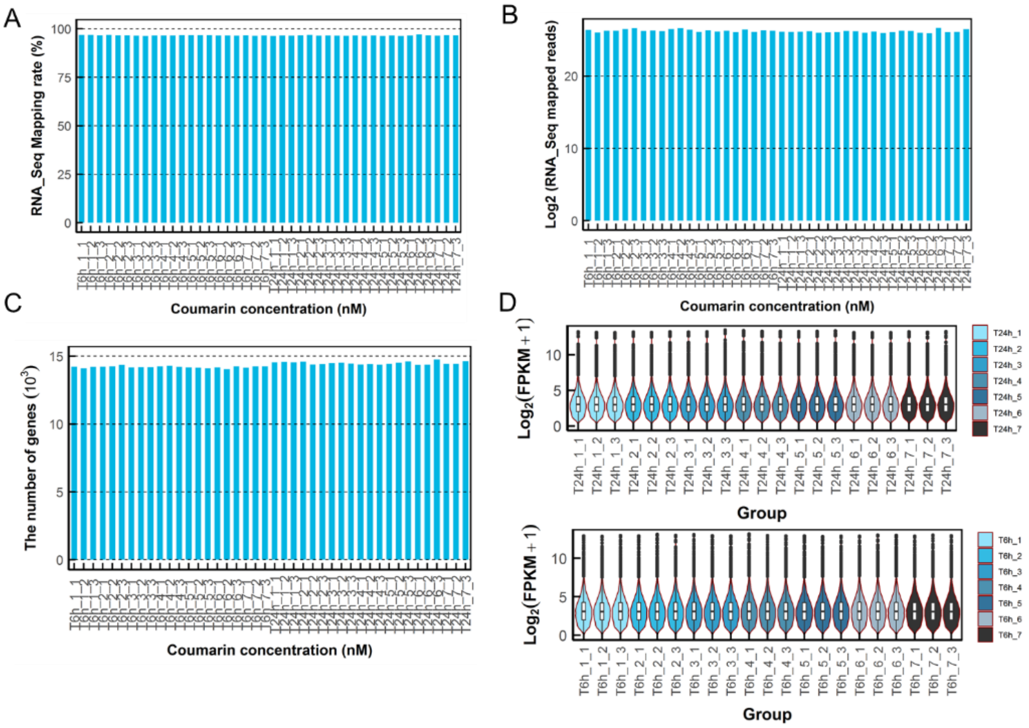


**Figure S1. Quality control for the transcriptomic study in Hep G2 cells treated with Coumarin.**

(A) Percentage RNA reads mapped to genome in each sample. (B) Number of mapped RNA reads in each sample. (C) Number of identified genes in each sample. (D) Violin and boxplots showing the statistics of FPKM in each sample. For RNA-seq, a FPKM cutoff value > 1.0 in at least one out of all samples was used to filter out false positives. Each sample is labelled as TAh-B-C, where T denotes transcriptome, A the hours (h) of cell treatment with coumarin, B the coumarin concentration numeric code as indicted in Figure 1B, and C the replicate number. Except for T, same letter denotations apply to other figures.

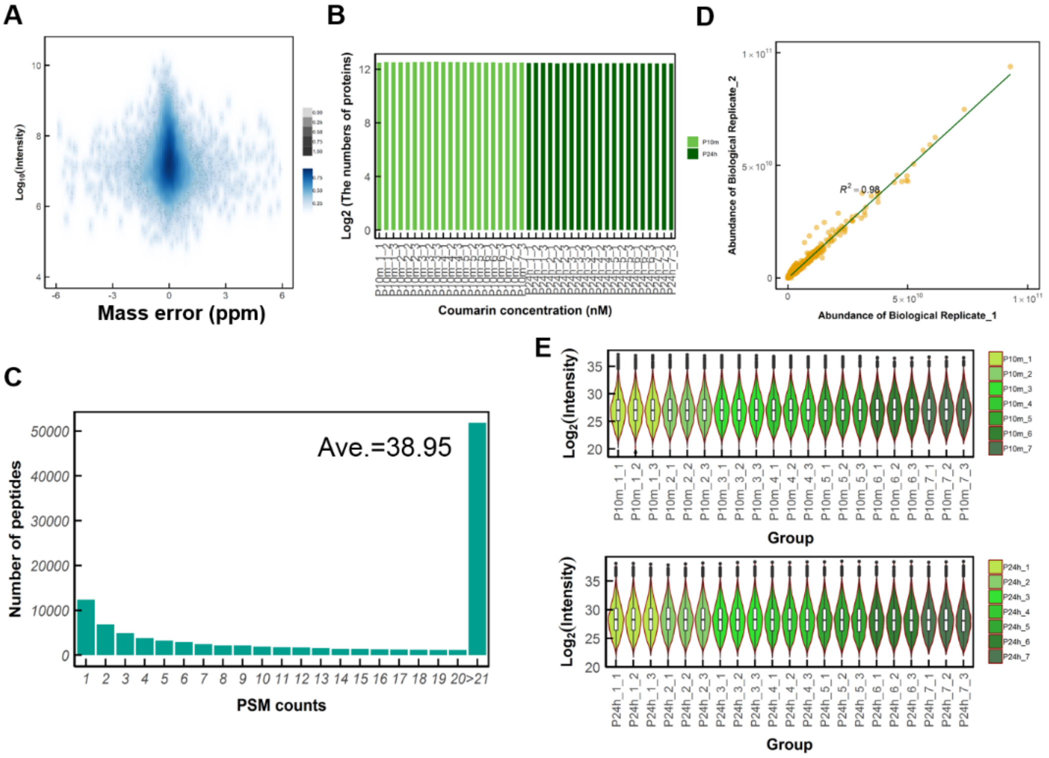


**Figure S2. Quality control for the proteomic study in Hep G2 cells treated with Coumarin.**

(A) Distribution of mass error of each matched spectrum for identified peptides. (B) Number of proteins identified in each sample. (C) Distribution of number of MS/MS spectrum per peptide. (D) Pearson correlation analysis of protein abundance (LFQ) between two control biological replicates. (E) Violin and boxplots showing the statistics of intensity in each sample. Peptides with intensities detected in at least 4 out of 21 samples were kept for further analysis and used to impute the missing intensities in remaining samples using the normal distribution. The letter P in sample label denotes proteome.

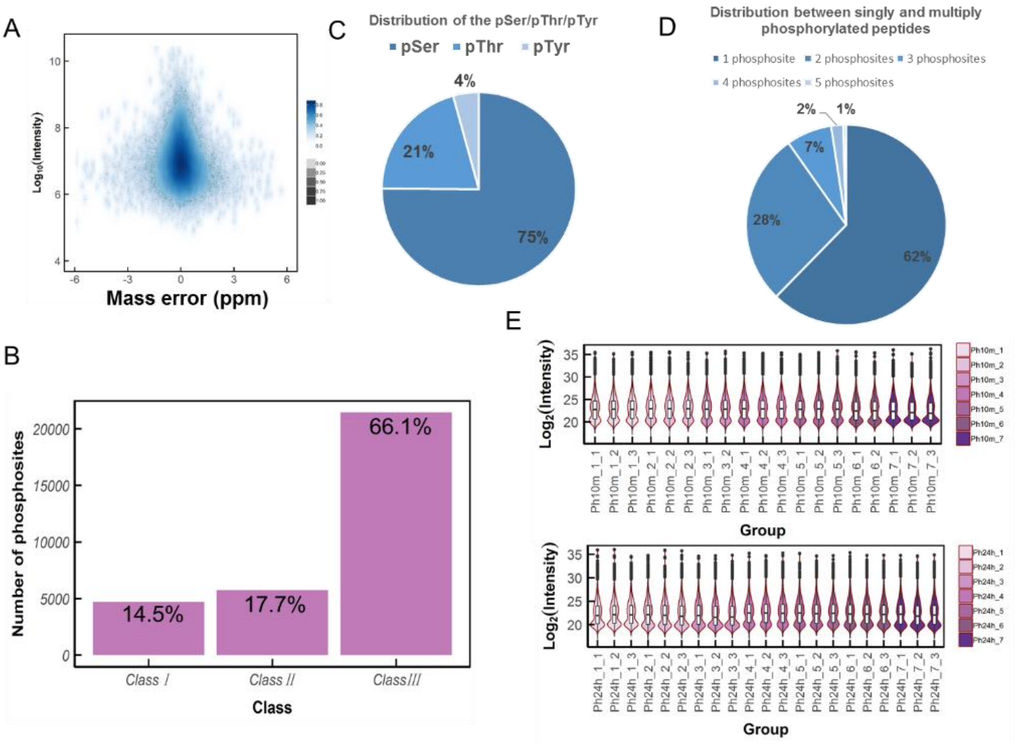


**Figure S3. Quality control for the phospho-proteomic study in Hep G2 cells treated with Coumarin.**

(A) Distribution of mass error of each matched spectrum for identified phosphopeptide. (B) Distributions of localization confidence for identified phosphosites. Classes 1, 2, and 3 represent phosphosites with probability scores between 0.25-0.5 (low confidence), 0.5-0.75 (medium confidence), and greater than 0.75 (high confidence), respectively. (C) Distribution of three types of phosphopeptides (i.e. pSer, pThr and pTyr). (D) Distribution of singly and multiply phosphorylated peptides. (E) Violin and boxplots showing the statistics of intensity in each sample. Phosphosites with intensities detected in at least 4 out of 21 samples were kept for further analysis and used to impute the missing intensities in remaining samples using the normal distribution in Perseus software. The letter Ph in sample label denotes phosphoproteome.

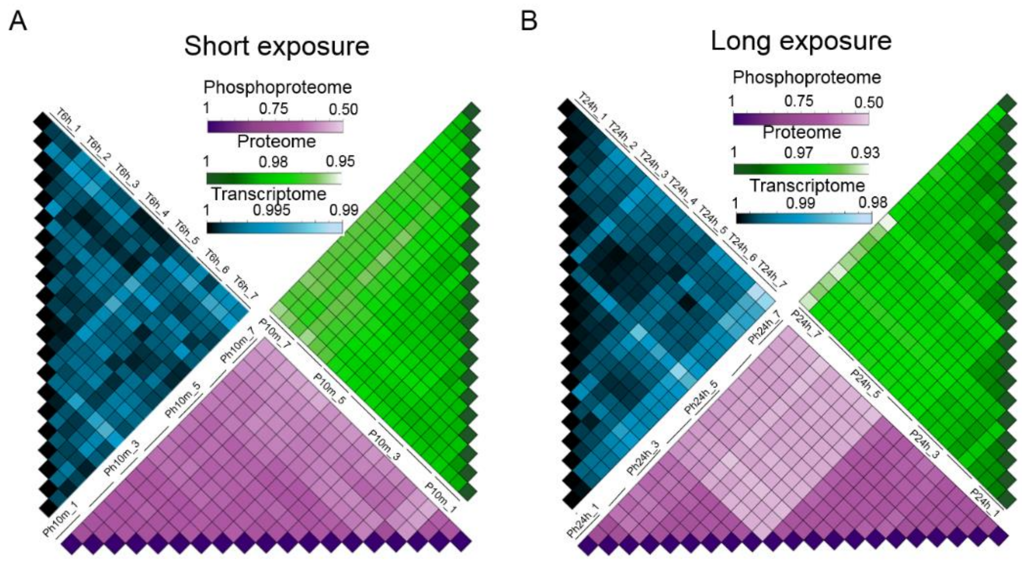


**Figure S4. Pair-wise Pearson correlation analysis of phosphoproteomic, proteomic, and transcriptomic samples of Hep G2 cells treated with Coumarin.** Heatmap representation of Pearson correlation coefficients between all replicate samples for (A) short exposure (10 min or 6 hr) or (B) long exposure (24 hr). Blue: transcriptome; green: proteome; purple: phosphoproteome.

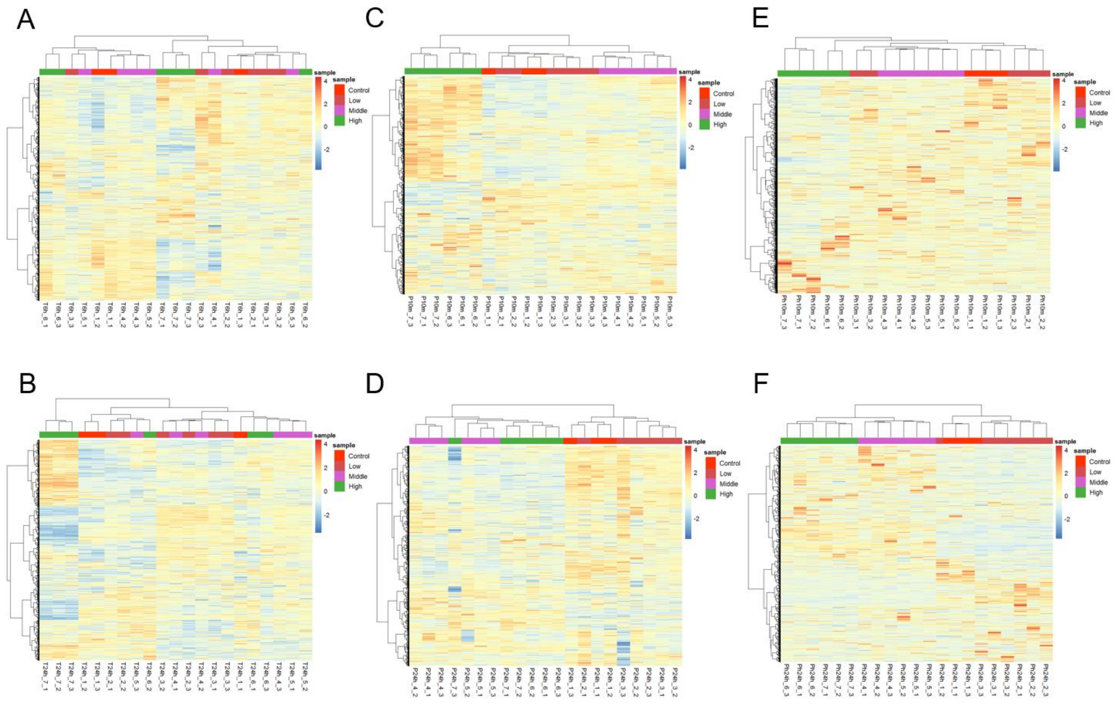


**Figure S5. Hierarchical clustering of the transcriptome, proteome, and phosphoproteome of HepG2 cells treated with Coumarin.**

(A, C & E) Unsupervised hierarchical clustering of transcriptomic (A), proteomic (C), and phosphoproteomic (E) data for short exposure (10 min or 6 hr). (B, D & F) Unsupervised hierarchical clustering of transcriptomic (B), proteomic (D), and phosphoproteomic (F) for long exposure (24 hr). The control, 0.001, 0.01 μM coumarin treatments are labelled red, the 0.1 and 1 μM treatment are labelled purple, and the 10 and 100 μM treatment are labelled green.

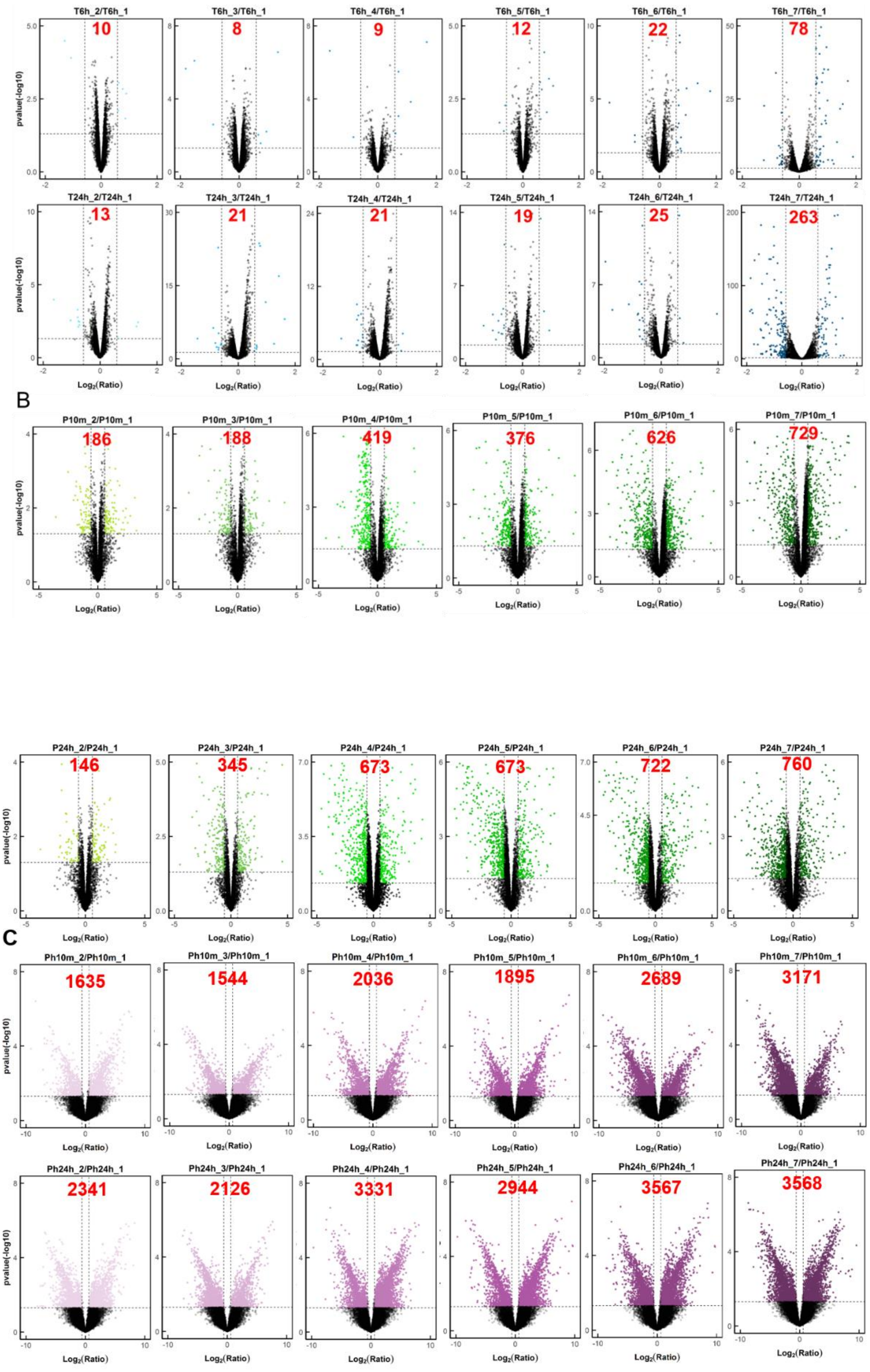


**Figure S6. Volcano plots of differentially expressed features in the proteome,**

**phosphoproteome, and transcriptome of HepG2 cells treated with Coumarin.**

(A, B & C) Volcano plots of the log2 fold change vs. -log10 p-value for mRNA (A), protein (B) and phosphor-site (C) features at each coumarin concentration compared with control. p value < 0.05 and fold change > 1.5 or < 0.67 were used as cutoff (horizontal and vertical lines) values for significantly differentially altered genes, proteins and phospho-sites. Numbers of significantly altered features are indicated in each panel.

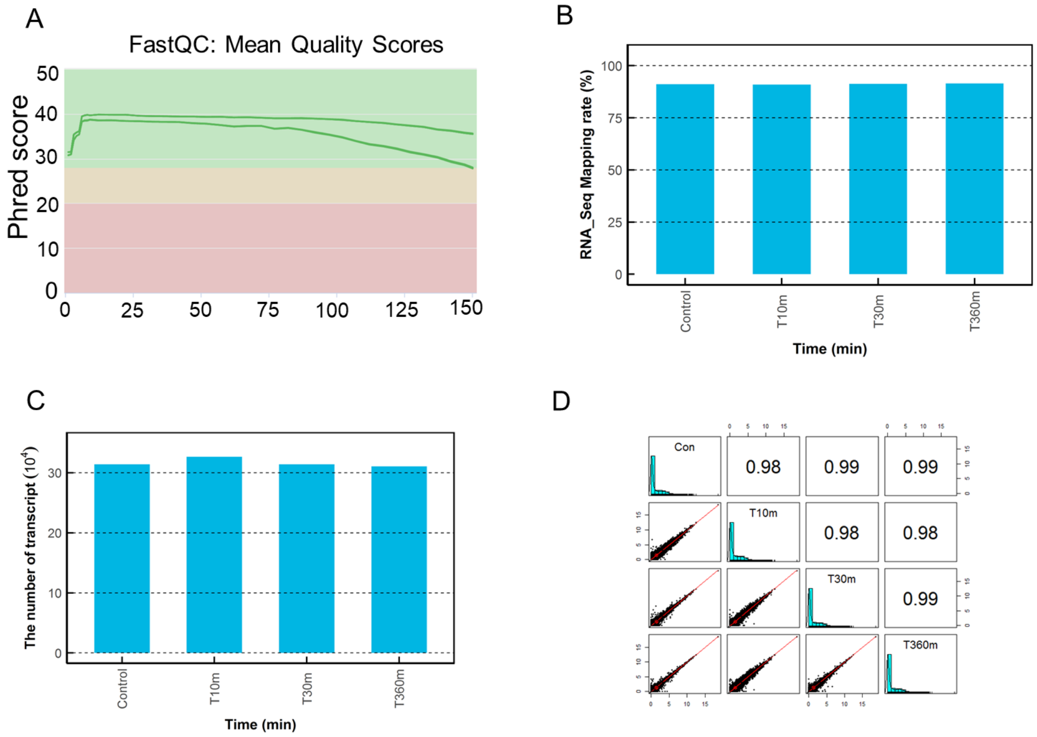


**Figure S7. Quality control for the transcriptomic study in AC16 cells treated with 10 nM doxorubicin for 10, 30, and 360 min.**

(A) Per sequence quality scores in FastQC. (B) The mapping rate of transcriptome in each sample. (C) The number of transcripts identified in each sample. (D) Pearson correlation analysis of gene abundance (FPKM) for each pair of samples.

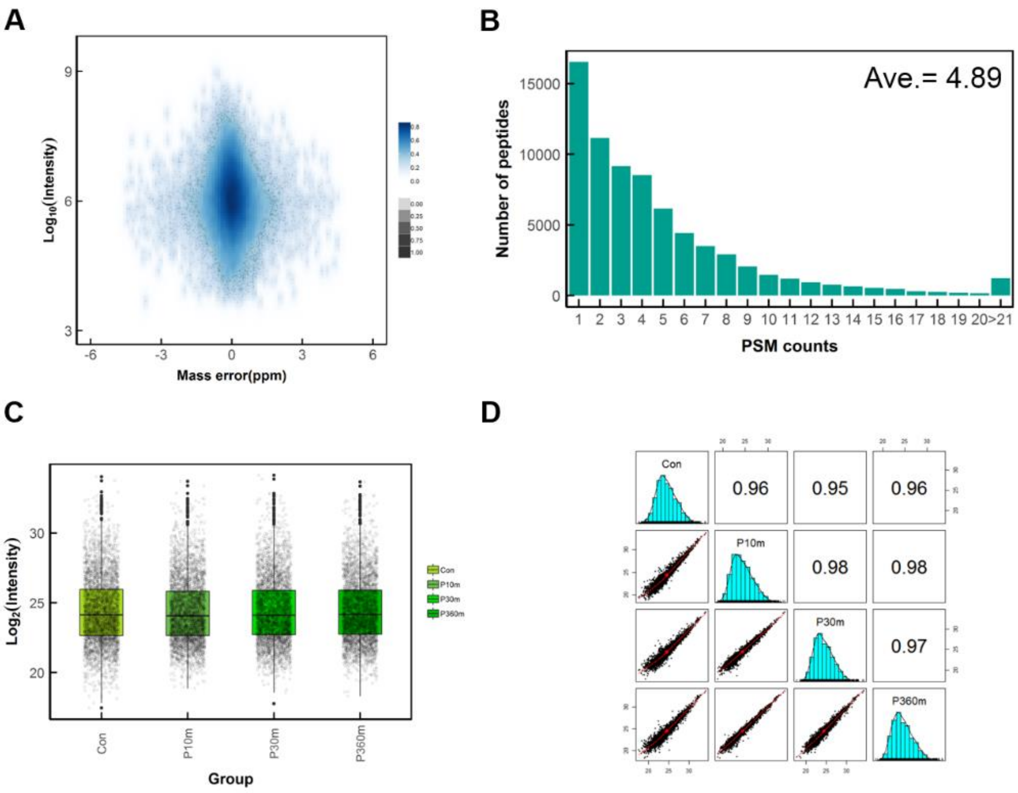


**Figure S8. Quality control for the proteomic study in AC16 cells treated with 10 nM doxorubicin for 10, 30, and 360 min.**

(A) Distribution of mass error of each matched spectrum for the identified peptides. (B) Distribution of number of MS/MS spectrum per peptide. (C) Identification and quantitation of proteins in each sample. (D) Pearson correlation analysis of proteins abundance (LFQ) for each pair of groups.

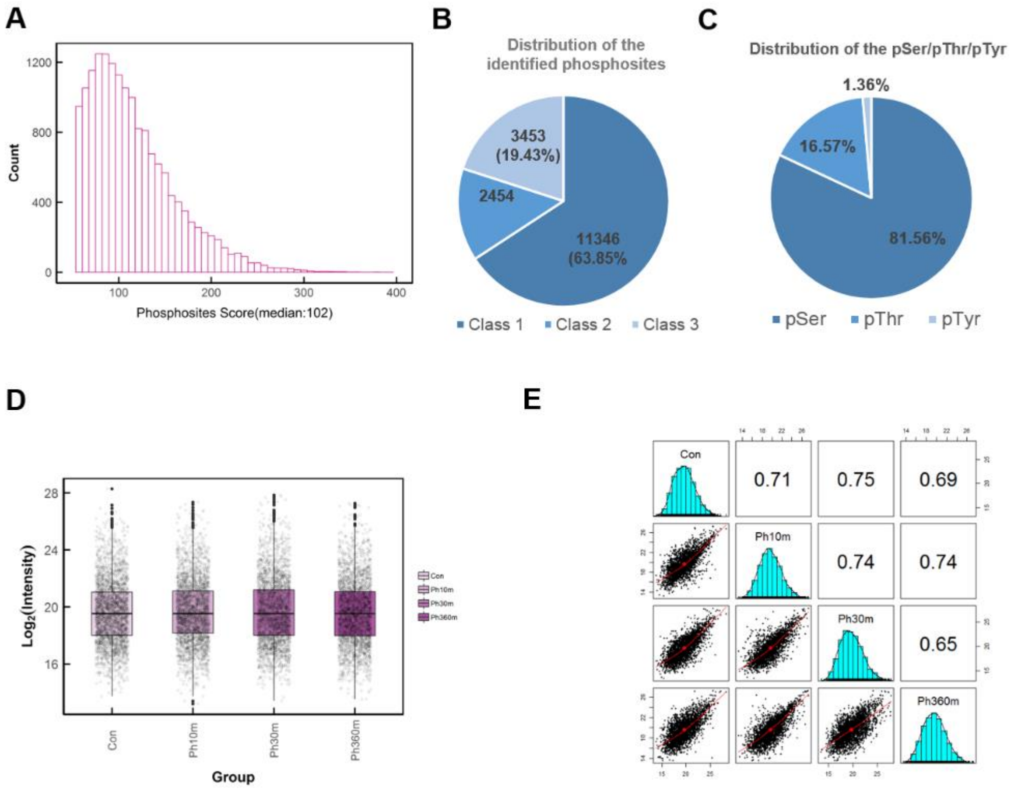


**Figure S9. Quality control for the phosphor-proteomic study in AC16 cells treated with 10 nM doxorubicin for 10, 30, and 360 min.**

(A) Distribution of phosphosites score. (B) Distributions of localization confidence for identified phosphosites. Different classes represent phosphosites with probability scores between 0.25-0.5 (low confidence), 0.5-0.75 (medium confidence), and greater than 0.75 (high confidence), respectively. (C) Distribution of three types of phosphosites (i.e. pSer, pThr and pTyr). (D) Identification and quantitation of phosphosites in each sample. (E) The Pearson correlation analysis of phosphosite abundance for each pair of groups.

### Supplementary Tables

Table S1. The details of AC50 values in coumarin treated HepG2 cells from literature.

Table 2. Transcripts identified in HepG2 cells treated with different coumarin concentration for 6 or 24 h.

Table S3. Proteins identified in HepG2 cells treated with different coumarin concentrations for 10 min.

Table S4. Proteins identified in HepG2 cells treated with different coumarin concentrations for 24 h.

Table S5. Phosphosites identified in HepG2 cells treated with different coumarin concentrations for 10 min.

Table S6. Phosphosites identified in HepG2 cells treated with different coumarin concentrations for 24 h.

Table S7. Transcripts identified in AC16 cells treated with 10 nM doxorubicin for 10, 30, or 360 min.

Table S8. Proteins identified in AC16 cells treated with 10 nM doxorubicin for 10, 30, or 360 min.

Table S9. Phosphosites identified in AC16 cells treated with 10 nM doxorubicin for 10, 30, or 360 min.

Table S10. The number of 194 phosphosites(cluster6) identified from AC16 cells in different time with doxorubicin treatment.

Table S11. The number of 157 proteins (match 195 phosphosites) identified from AC16 cells in different time with doxorubicin treatment.

Table S12. HMOX1 protein identified in AC16 cells treated with 10 nM doxorubicin for 10, 30, or 360 min.

Table S13. TSC2 phosphosites identified in AC16 cells treated with 10 nM doxorubicin for 10, 30, or 360 min.

Table S14. SQSTM1 phosphosites identified in AC16 cells treated with 10 nM doxorubicin for 10, 30, or 360 min.

Table S15. Parameter values of the mathematical model.

Table S16. Ordinary differential equations (ODEs) of the mathematical model.
