## Supplementary material for "The phosphoproteome is a first responder in tiered cellular adaptation to chemical stress followed by proteomics and transcriptomics alteration": Table S15

**Table S15. Parameter values of the mathematical model.**

| **Parameter** | **Value** | **Unit** | **Note** |
| --- | --- | --- | --- |
| $k_{0}$ | 0.1 | Mass/L/S | Basal production rate of controlled state Y |
| $k_{1}$ | 0.1 | 1/S | Rate constant of stressor S-induced production of Y |
| $k_{2}$ | 0.1 | 1/(Mass/L)/S | Rate constant of active stress protein Ga-catalyzed degradation of Y |
| $k_{3}$ | ln(2)/(30*60) | 1/S | Rate constant of Y-activated synthesis of transcription factor T |
| $k_{4}$ | ln(2)/(30*60) | 1/S | Rate constant of degradation of T, corresponding to a half-life of 30 min |
| $k_{5}$ | 5*ln(2)/(3*3600) | Mass/L/S | Rate constant of T-induced transcription of M |
| $k_{6}$ | ln(2)/(3*3600) | 1/S | Rate constant of degradation of stress mRNA, corresponding to a half-life of 3 h |
| $k_{7}$ | 6*ln(2)/(6*3600) | 1/S | Rate constant of M-stimulated translation of inactive stress protein Gi |
| $k_{8}$ | ln(2)/(6*3600) | 1/S | Rate constant of degradation of inactive stress protein Gi, corresponding to a half-life of 6 h |
| $k_{9}$ | 0.002 | 1/S | Catalytic rate constant of active kinase Aa-catalyzed phosphorylation (activation) of Gi to Ga |
| $k_{10}$ | 0.002 | 1/S | Vmax of dephosphorylation (deactivation) of Ga to Gi |
| $k_{11}$ | ln(2)/(6*3600) | 1/S | Rate constant of degradation of Ga, corresponding to a half-life of 6 h |
| $k_{12}$ | 0.01 | 1/S | Rate constant of Y-induced activation of Ai to Aa |
| $k_{13}$ | 0.01 | 1/S | Rate constant of deactivation of Aa to Ai |
| $k_{14}$ | 140*ln(2)/(10*3600) | Mass/L/S | Rate constant of Y-inhibited synthesis of cell function protein CFP |
| $k_{15}$ | ln(2)/(10*3600) | 1/S | Rate constant of degradation of CFP, corresponding to a half-life of 10 h |
| $k_{50}$ | 0.5*ln(2)/(3*3600) | Mass/L/S | Basal transcription rate of stress mRNA (M) |
| $n_{5}$ | 5 | - | Hill coefficient of T-induced transcription of M |
| $n_{14}$ | 5 | - | Hill coefficient of Y-inhibited synthesis of CFP |
| $J_{5}$ | 10^(log10(9)/5) | Mass/L | Affinity constant of T-induced transcription of M |
| $J_{9}$ | 0.1 | Mass/L | Michaelis constant of Aa-catalyzed phosphorylation (activation) of Gi to Ga |
| $J_{10}$ | 0.1 | Mass/L | Michaelis constant of dephosphorylation (deactivation) of Ga to Gi |
| $J_{12}$ | 0.1 | Mass/L | Michaelis constant of Y-induced activation of Ai to Aa |
| $J_{13}$ | 0.1 | Mass/L | Michaelis constant of deactivation of Aa to Ai |
| $J_{14}$ | 1.2 | Mass/L | Affinity constant of Y-inhibited synthesis of CFP |
| ${tau}_{14}$ | 1800 | S | Time delay in Y-inhibited synthesis of CFP |
| $A_{tot}$ | 10 | Mass/L | Total kinase |
| S | varied | Mass/L | Chemical stressor |
