## Supplementary material for "The phosphoproteome is a first responder in tiered cellular adaptation to chemical stress followed by proteomics and transcriptomics alteration": Table S16

**Table S16. Ordinary differential equations (ODEs) of the mathematical model.**

| **Variable** | **ODE** | **Steady-state level** |
| --- | --- | --- |
| Y  (Controlled state) | dY/dt = k0 + k1*S - k2*Ga*Y |  |
| T  (Transcription factor) | dT/dt = k3*Y - k4*T |  |
| M  (mRNA) | dM/dt = k50 + k5*T^n5/(J5^n5+T^n5) - k6*M |  |
| Gi  (Inactive stress protein) | dGi/dt = k7*M - k8*Gi - k9*Aa*Gi/(J9+Gi) + k10*Ga/(J10+Ga) |  |
| Ga  (Active stress protein) | dGa/dt = k9*Aa*Gi/(J9+Gi) - k10*Ga/(J10+Ga) - k11*Ga |  |
| Aa  (Active kinase) | dAa/dt = k12*Y*(Atot - Aa)/(J12+Atot - Aa) - k13*Aa/(J13+Aa) |  |
| CFP  (Cell function protein) | dCFP/dt = k14*J14^n14/(J14^n14+(DELAY(Y, tau14))^n14) - k15*CFP |  |
